# FERONIA’s sensing of cell wall pectin activates ROP GTPase signaling in *Arabidopsis*

**DOI:** 10.1101/269647

**Authors:** Wenwei Lin, Wenxin Tang, Charles T. Anderson, Zhenbiao Yang

## Abstract

Plant cells need to monitor the cell wall dynamic to control the wall homeostasis required for a myriad of processes in plants, but the mechanisms underpinning cell wall sensing and signaling in regulating these processes remain largely elusive. Here, we demonstrate that receptor-like kinase FERONIA senses the cell wall pectin polymer to directly activate the ROP6 GTPase signaling pathway that regulates the formation of the cell shape in the *Arabidopsis* leaf epidermis. The extracellular malectin domain of FER directly interacts with de-methylesterified pectin *in vivo* and *in vitro*. Both loss-of-FER mutations and defects in the pectin biosynthesis and de-methylesterification caused changes in pavement cell shape and ROP6 signaling. FER is required for the activation of ROP6 by de-methylesterified pectin, and physically and genetically interacts with the ROP6 activator, RopGEF14. Thus, our findings elucidate a cell wall sensing and signaling mechanism that connects the cell wall to cellular morphogenesis via the cell surface receptor FER.

## INTRODUCTION

Mounting evidence suggests that cell wall polymers provide signals to regulate a large number of plant processes and that plant cells can monitor cell wall dynamics or damages ^1" rid="c5">5^. For instance, cell wall biosynthesis and cell wall remodeling-related genes are reprogramed in various cell wall-related mutants^6,7^ or in plants treated with cellulose synthesis inhibitor ^8^ or pectin fragments ^9^. Upon the interferences with cell wall status, various compensatory wall responses were induced, including ectopic deposition of lignin and callose ^10" rid="c12">12^, altered pectin modification status^8,13,14^, reactive oxygen species (ROS) production ^11^, and an elevation in jasmonic acid and ethylene production^11,15,16^. Furthermore, cell wall dynamic plays a regulatory role in the regulation of complex cell shapes ^7^. However, the mechanisms for sensing and transducing the wall signals remain largely mysterious. The elucidation of these mechanisms is solely needed to understand how cell expansion and shape changes are coordinated between neighboring cells in a growing tissue or organ.

The spatiotemporal pattern of cell walls (and as such their compositions and structure) is critical for cell expansion and shape formation, because of high cellular turgor pressure in plants^17,18^. Precise coordination and communication via the dynamic cell wall between adjacent cells to regulate these cellular processes are required for organ differentiation, growth, and morphogenesis, and yet the underlying mechanisms remain elusive. The jigsaw puzzle-shaped pavement cells (PCs) of the leaf epidermis serve as an exciting model to investigate the mechanisms for cell-cell coordination of cell shape in a multicellular system ^19^. PCs form interdigitated lobes and indentations that are tightly coordinated between the adjacent cells. This process is regulated by two antagonistic RAC/ROP GTPase signaling pathways in an auxin-dependent manner ^20^. Locally activated ROP2/ROP4 signaling promotes the outgrowth to form the lobe region, whereas ROP6 at the indenting region promotes the ordering of cortical microtubules (MTs), which restrict radial or lateral cell expansion ^21^.

Mechanical stress from lobe outgrowth has been implicated in the indentation reinforcement by promoting cortical MT ordering ^22^, hinting at a potential regulatory role of the cell wall in this process. This is further supported by a recent study suggesting that the cell wall compositions and mechanical heterogeneities across and along anticlinal cell walls contribute to the regulation of PC interdigitation^7^. The identification of cell wall sensors is needed to understand the mechanisms by which the cell wall modulates various plant processes such as PC interdigitation. A large number of cell surface receptors, such as Wall-associated kinases (WAKs), *Catharanthus roseus* RLK1 (CrRLK1)-like family, PR5-like receptor kinase/thaumatin family, LysM family, L-type lectin RLKs, proline-rich extension like receptor kinase (PERK) family, leucine-rich repeat extensins (LRXs), FEI1/2, LRR-RK MALE DISCOVERER 1-INTERACTING RECEPTOR LIKE KINASE 2/LEUCINE-RICH REPEAT KINASE FAMILY PROTEIN INDUCED BY SALT STRESS (MIK2/LRR-KISS) and LRR receptor-like protein 44 (RLP44) have been proposed to be potential wall sensors^1,5,23,24 25" rid="c34">34^, but none of them have been clearly demonstrated to sense and transduce cell wall polymer signals. Here we report that the FERONIA cell surface receptor senses and transmits cell wall pectin to activate the ROP signaling pathway to modulate the formation of the puzzle-piece cell shape in *Arabidopsis* PCs.

## RESULTS

### The FERONIA (FER) cell surface receptor is associated with the cell wall

Plants have evolved a large number of cell surface receptor-like kinases (RLKs) to sense various extracellular signals ^35^. To identify RLKs that might be involved in the lobe-indentation coordination during PC interdigitation, we carried out a genetic screen of T-DNA insertion *rlk* mutants for altered PC morphogenesis and isolated *fer-4*, a mutant of *FERONIA* ^36^, which exhibited a severe defect in PC shape (Fig. 1a, b). Besides root hair growth and development, FER was suggested to be involved in cell polarization, which is important for the formation of the interdigitated shape in *Arabidopsis* leaf epidermal PCs ^37^. To further evaluate the role of FER in PC morphogenesis, we characterized another (*fer* allele with an independent T-DNA insertion (*fer-2* and *fer-5*, Supplementary Fig. 1a)^38,39^ and found that both alleles showed similar defects in PC interdigitation. Both *fer* mutants displayed significantly wider indentations than wild-type (WT) (Fig. 1a, b, and Supplementary Fig. 1b-d), reminiscent of a defect in the ROP6 signaling pathway ^20^. To further confirm that the phenotypes observed in the *fer* mutants were due to the mutation in *FER*, we complemented the *fer-4* mutant with *FER* cDNA fused with yellow fluorescent protein (*YFP*) under the control of its native promoter. The *FER-YFP* transgene completely rescued the *fer-4* PC shape defect and restored vegetative growth to wild-type levels (Fig. 1a, b, and Supplementary Fig. 1e). Together, the data indicate that FER is required for PC morphogenesis.

**Figure 1.**
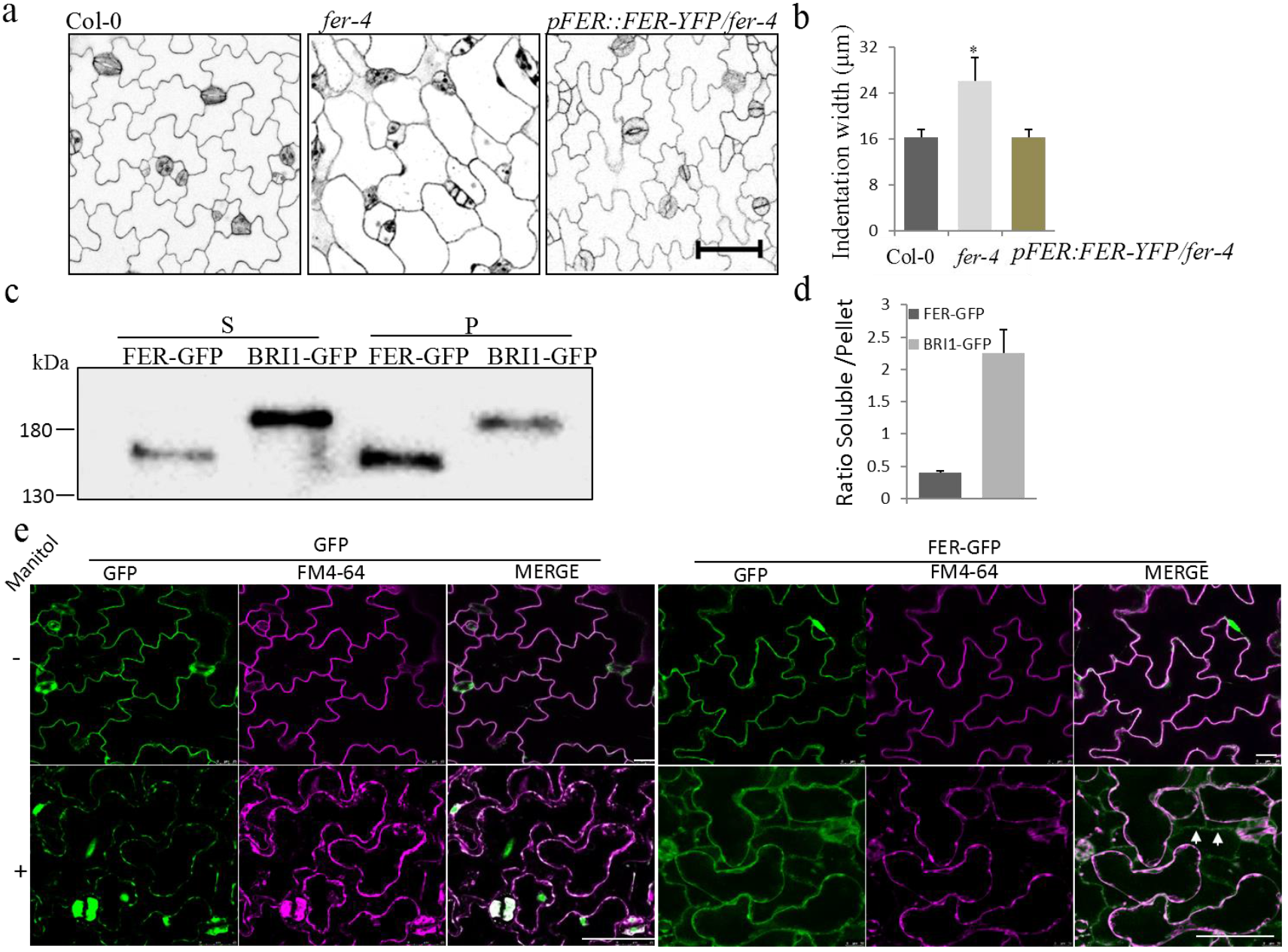
FER is associated with the cell wall and required for *Arabidopsis* epidermal pavement cells (PCs) morphogenesis. **a and b**, PC phenotypes of 2 day-after-germination (DAG) wild-type (Col-0), *fer-4* and complementary line *pFER::FER-YFP/fer-4* (a). The degree of PC interdigitation was determined by the width of indentation (b). Data was generated from the measurement of at least 20 cells collected from 5 different cotyledons from 5 individual seedlings. Asterisk indicates a significant difference with p⩽0.05 (Student’s *t*-test) between wild-type and the *fer-4* mutant. Data are represented as mean ± SE. Scale bar, 50 μm. **c and d**, FER-GFP proteins are enriched in the insoluble fraction. Soluble (S) and insoluble (P) proteins were fractionated and analyzed by Western blot (WB) with α-GFP antibody (c) and the ratio of two fractions was quantified (d). The plasma membrane-localized RLK *BRI1-GFP* transgenic plant was used as a control. The data are shown as mean ± SE of three repeats. **e**, The cell wall localization of FER-GFP. (*Top* panels) GFP fluorescence was along with the cell surface of PCs before plasmolysis (−). During plasmolysis (+) GFP fluorescence localized with the cytoplasm (*Bottom* left panel) and plasma membrane indicated by overlapping signal with FM4-64 (Merge). A portion of the FER-GFP signal retreated with the cytoplasm localized on the plasma membrane indicated by overlapping signal with FM4-64 (Merge) and another portion of the signal remained on the cell surface (*Bottom* right panel). Arrows indicate the cell wall residue FER-GFP signal. Scale bar, 50 μm.

FER belongs to the *Arabidopsis* subfamily of *Catharanthus roseus* RLK1-like kinases (CrRLK1Ls) with 17 members all containing a malectin-like domain predicted to bind carbohydrates (Supplementary Fig. 1a)^3^. In *Arabidopsis*, *FER* was initially found to control pollen tube perception in the ovule^36^ and later shown to regulate a wide range of growth and developmental processes and responses to the environment^38" rid="c46">46^. Interestingly several CrRLK1L members play important roles in controlling processes linked to the properties of the cell wall, including cell expansion, polarized growth, sperm release from pollen tubes, pollen tube integrity maintenance, and defense responses^24,36,40,41,47" rid="c49">49^. In *Arabidopsis*, FER and its relatives, ANUXR1, ANUXR2, BUPS1, and BUPS2 are involved in maintaining cell wall integrity and to regulate cell growth through the recognition of a group of small peptide ligands that derived from cells named Rapid Alkanilization Factors (RALFs)^24,48,50^. Another CrRLKL1 family member, THESUS1(THE1), identified as a putative cell-wall integrity sensor, mediates the responses to the perturbation of cellulose synthesis, likely via binding to the cell wall^3,30^. Therefore, we speculated that FER might have a role as a cell wall sensor to monitor the dynamics of the cell wall during PCs morphogenesis, and assessed whether FER is associated with the cell wall. We generated transgenic plants expressing FER-green fluorescent protein (GFP) fusion driven by its native promoter. BRI1, a known plasma membrane-localized protein, was used as a control ^51^. Soluble (supernatant) and insoluble (pellet) proteins were fractionated from FER-GFP and BRI1-GFP transgenic plants by using low-speed centrifugation. As expected, non-wall-associated BRI1-GFP was enriched in the soluble fraction (Fig. 1c, d). In contrast, FER-GFP was enriched in the insoluble pellet fraction (Fig. 1c, d). As shown for wall-associated kinase proteins (WAKs)^28,52^, the extraction of FER-GFP from the insoluble pellet fraction required boiling in the presence of 1% (w/v) SDS and 50 mM dithiothreitol (DTT) (Supplementary Fig. 1f), indicating that FER is tightly associated with the cell wall.

To further verify the cell wall association of FER, cotyledon PCs from transgenic seedlings expressing FER-GFP fusion or GFP alone were plasmolyzed (Fig. 1e). Before plasmolysis, the GFP control exhibited a typical free-GFP localization pattern in plant cells with a large central vacuole; the signals were largely diffused along the cell border (cell walls stained by Propidium Iodide, PI) (Supplementary Fig. 1g). After plasmolysis, the GFP signals retreated with the plasma membrane (plasma membrane stained by FM4-64) that was detached from the cell walls (Fig. 1e, Supplementary Fig. 1g, the lower row of panels). By contrast, FER-GFP was mainly found along the plasma membrane, largely overlapping with the cell wall PI staining (Supplementary Fig. 1g). After plasmolysis, a portion of the FER-GFP fluorescence retreated with the plasma membrane (white arrows), while predominant signal remained at the cell border (Fig. 1e, Supplementary Fig. 1g, the lower row of panels). These results are consistent with the high enrichment of the FER-GFP protein in the cell wall fraction, suggesting that FER is a plasma membrane RLK that is tightly associated with the cell wall.

**Figure 2.**
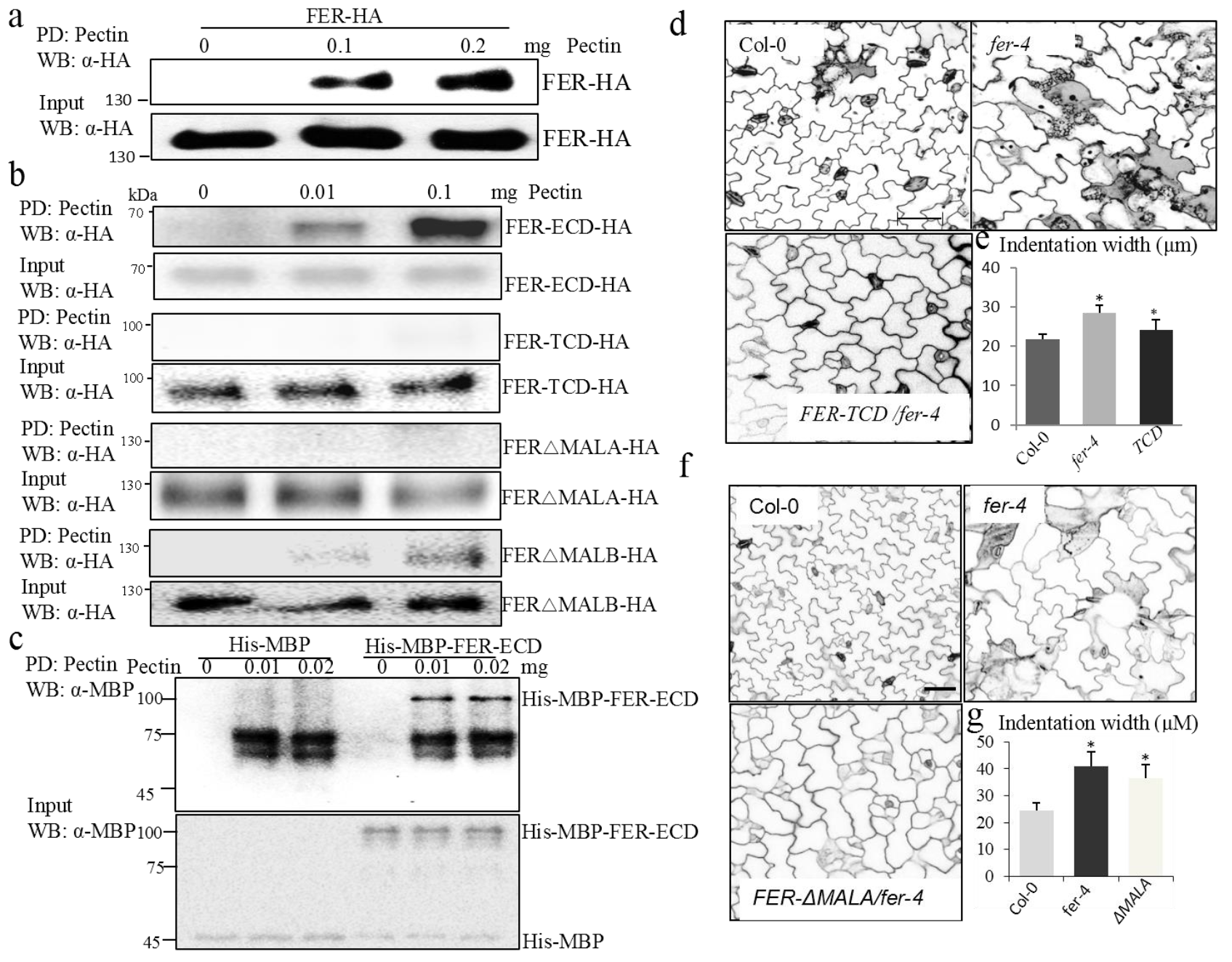
FER binds pectin through the MALA domain to regulate PC morphogenesis. **a**, FER was associated with pectin *in vitro*. FER-HA was transiently expressed in *Arabidopsis* protoplasts. Pull-down (PD) was carried out with pectin and the proteins were determined by Western blot (WB) with α-HA antibody. *Top* shows that FER-HA pull-down by pectin (PD: pectin; WB: α-HA). *Bottom* shows the expression of FER-HA proteins (WB: α-HA). **b**, Association of truncated FER with pectin *in vitro*. Different specific deletion domains of FER-HA were expressed in *Arabidopsis* protoplasts, the PD assays were carried out with pectin. *Top* shows that truncated FER-HA was pulled-down by pectin (PD: pectin; WB: α-HA). *Bottom* shows the expression of FER-HA truncated proteins (WB: α-HA). **c**, FER-ECD directly bound to pectin *in vitro*. His-MBP-FER-ECD was purified from *E. coli*. Pull down was carried out as above and the proteins were determined by WB with α-MBP antibody. His-MBP-FER-ECD pull-down by pectin (*Top*). The input His-MBP and His-MBP-FER-ECD proteins (*Bottom*). **d**, PC phenotypes in wild-type, *fer-4* and the complementary line *35S::YFP-FER-TCD/fer-4*. Scale bar, 20 μm. **e**, The degree of 3 DAG PC interdigitation. Asterisk indicates the average indentation widths that were significantly different (p≤0.05, Student’s *t*-test) between wild-type and the *fer-4* mutant, wild-type, and *35S::YFP-FER-TCD* complementation line, respectively. **f**, PC phenotypes in wild-type, *fer-4* and the complementary line *35S::YFP-FER-ΔMALA/fer-4. Scale bar, 50* μm. **g**, The degree of 6 DAG PC interdigitation. Asterisk indicates the average indentation widths that were significantly different (p≤0.05, Student’s *t*-test) between wild-type and *fer-4* mutants, wild and the *35S::YFP-FER-ΔMALA* complementation line, respectively.

### The extracellular malectin A domain is responsible for FER’s association with pectin and is required for FER’s function in PCs

We next determined whether FER is associated with a specific cell wall component using a semi-*in vitro* pull-down assay. The FER protein fused to an HA tag was expressed in *Arabidopsis* leaf protoplasts, and proteins isolated from the protoplasts were incubated with different cell wall components. Insoluble cellulose and hemicellulose (xylan) were pelleted by centrifugation, but FER-HA was not detected in either the cellulose or xylan pellets (Supplementary Fig. 2a). In contrast, FER-HA was detected in the pellet of pectin extracted from apple (50-75% degree esterification) in a concentration-dependent manner (Fig. 2a), indicating an interaction between FER and pectin. We then dissected the domains of FER responsible for binding pectin (Supplementary Fig. 2b). FER-ECD-HA (where the intracellular domain was deleted) and FER△MALB-HA (deletion of the extracellular MALB domain) were pulled down by pectin, but FER-TCD-HA (deletion of the entire extracellular domains) and FER△MALA-HA (deletion of the extracellular MALA domain) were not (Fig. 2b). Importantly, we found that His-MBP-FER-ECD recombinant proteins, expressed and purified from *E. coli*, directly bound to pectin *in vitro* (Fig. 2c). This suggests that glycosylation is not required for FER’s function in binding to pectin. Furthermore, in agreement with the finding that MALA domain is required for the interaction (Fig. 2b), we found that His-MBP-FER-MALA domain recombinant proteins, expressed and purified from *E. coli*, directly bound pectin *in vitro* (Supplementary Fig. 2c). Thus, these results demonstrate that FER physically binds to pectin through the extracellular MALA domain.

We then determined whether FER’s extracellular MALA domain is also required for the in vivo association with the cell wall and function. We generated *35S::YFP-FER-TCD* transgenic plants and detected the protein levels in both soluble and insoluble protein fraction using a GFP antibody. In contrast to the full-length FER-GFP protein, which was enriched in the insoluble pellet fraction (Fig. 1c, d), YFP-FER-TCD was enriched in the soluble fraction (Supplementary Fig.2d). Furthermore, the fluorescent signal of YFP-FER-TCD in plasmolyzed PCs was absent from the cell walls (Supplementary Fig. 2e). Thus the cytoplasmic domain is not involved in FER’s association with the cell wall. Since the MALA domain bound pectin *in vitro* (Fig. 2b and Supplementary Fig. 2c), we further tested its requirement for cell wall association in the *35S::YFP-FER△MALA* transgenic *Arabidopsis* plants by using plasmolysis assays. In contrast to FER-GFP (Fig. 1e), the majority of YFP-FER△MALA regressed with the shrunk cytoplasm (Supplementary Fig. 2f), indicating that the MALA domain is critical for FER’s association with the cell wall. Finally, neither YFP-FER-TCD nor YFP-FER-△MALA was able to fully rescue the PC interdigitation defect in the *fer-4* knockout mutant (Fig. 2d-g), which was fully rescued by the full-length FER-YFP (Fig. 1a). Taken together, our results indicate that the extracellular MALA domain binds to cell wall pectin, allowing FER to associate with the cell wall, and regulates PC morphogenesis.

### Pectin methylation level is spatially regulated and important for PC shape

The generation of lobes and indentations in PCs results from the anisotropic growth of the cell wall and involves the preferential deposition of cellulose in the indentation region ^53^. As expected mutants defective in cellulose synthesis typically exhibit simpler PC shapes ^54 7^. The pectin composition and structure in the wall in much more complicated and has been proposed to have a regulatory role^3,55" rid="c60">60^. With the finding that FER’s association with pectin is critical for PC morphogenesis, we sought to determine the contribution of the cell wall pectin to the formation of PC shapes. In *Arabidopsis*, three types of pectic polysaccharides are found: homogalacturonan (HG), rhamnogalacturonan I (RG-I), and rhamnogalacturonan II (RG-II) ^61^. Mutants with reduced HG levels (*qua1-1* and *qua2-1*) exhibit decreased PC interdigitation^7^. We found that the *arad1arad2* mutant, which has reduced arabinans associated with rhamnogalacturonan I (RG-I), showed PC shape changes similar to those observed in the *fer* mutants with wider indentation necks (Fig. 3a, b). These results suggest that the specific composition and structure of pectic polysaccharides are critical for PC morphogenesis. This agrees with the importance of fine pectin structures in modulating other biological roles^7, 60^.

**Figure 3.**
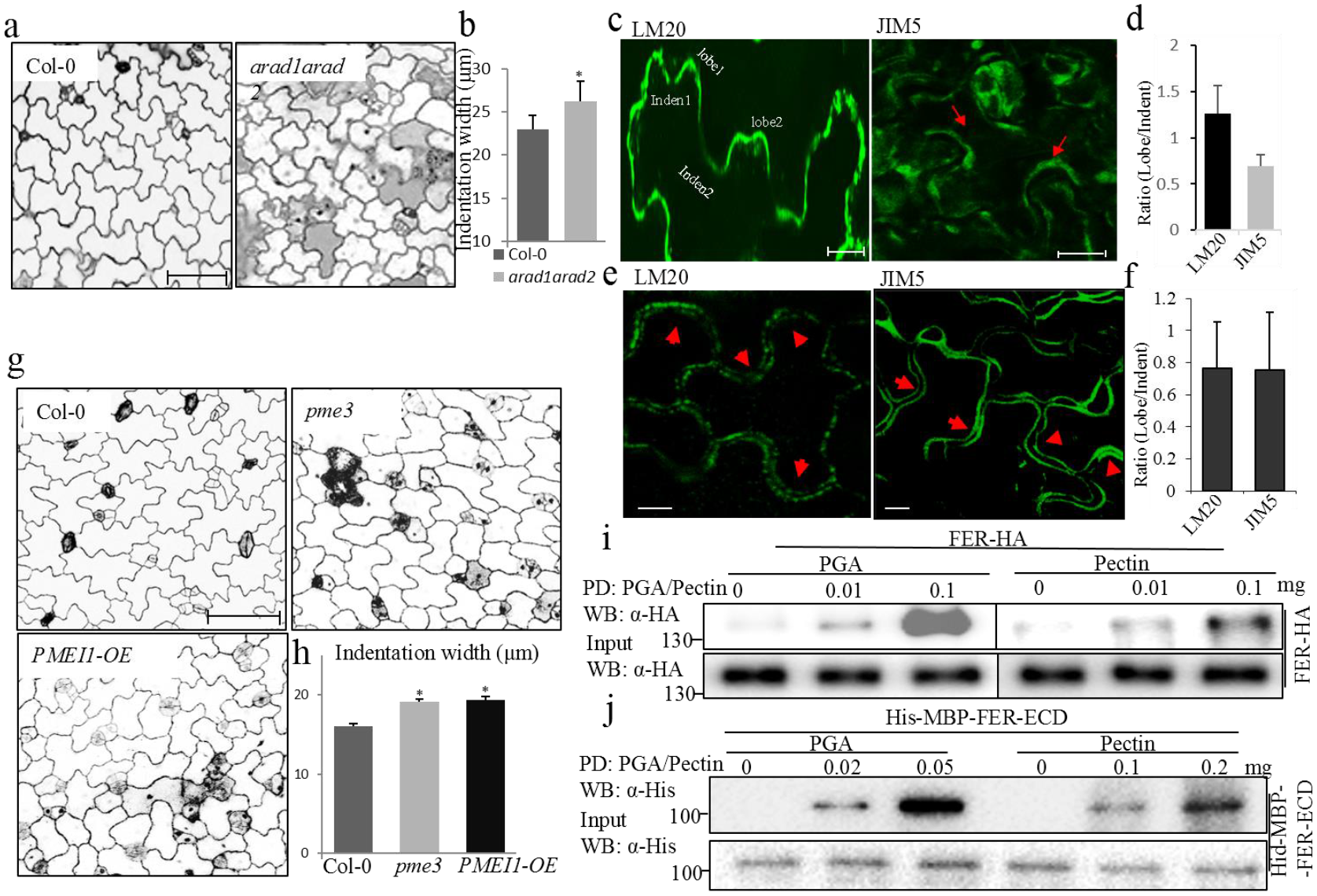
Pectin methylation levels modulate PC morphogenesis and pectin-FER interaction. **a**, Cotyledon PC phenotypes in wild-type and the *arad1arad2* mutant. Scale bar, 100 μm. **b**, The degree of 3 DAG PC interdigitation. Asterisk indicates a significant difference (p⩽0.05, Student’s *t*-test) between wild-type and the *arad1arad2* mutant. **c**, Expanding young PCs from 2 DAG cotyledons were immuno-stained for highly methylesterified or de-methylesterified pectin with LM20 (left) and JIM5 (right) antibodies, respectively. The lobe and indentation regions analyzed were indicated by label or arrows. Scale bar, 10 μm. **d**, Signal intensity ratios between lobe and indentation regions. The data are shown as mean ± SE of 20 cells. **e**, Mature PCs from 3 weeks-old seedlings **e**, (the third leaf pairs) were immunostained for highly methylesterified or de-methylesterified with LM20 (left) and JIM5 (right) antibodies, respectively. The lobe and indentation region analyzed were indicated by arrows. Scale bar, 10 μm. **f**, Signal intensity ratios between lobe and indentation regions. The data are shown as mean ± SE of 20 cells. **g**, Cotyledon PC phenotypes in wild-type, *pme3* and *PMEI1-OE* mutants. Scale bar, 100 μm. **h**, The degree of 2 DAG PC interdigitation. Asterisks indicate a significant difference (p⩽0.05, Student’ s *t*-test) between wild type and the mutants. **i**, FER was preferentially associated with de-methylesterified pectin (PGA) in protoplasts compared to highly methylesterified pectin (pectin). Pull-down was carried out with PGA and pectin as described above (Fig. 2a). (*Top*) FER-HA proteins from protoplasts were pull-down by PGA or pectin. (*Bottom*) The His-MBP-FER-ECD proteins were determined by WB as input control. **j**, FER-ECD preferentially bound de-methylesterified pectin (PGA) *in vitro* compared to highly methylesterified pectin (pectin). (*Top*) His-MBP tagged FER-ECD recombinant proteins purified from *E. coli* were pulled down by PGA or pectin and detected by α-His antibody. (*Bottom*) The His-MBP-FER-ECD proteins were determined by western blotting (WB) as input control.

Highly methylesterified pectic polysaccharides are synthesized in the Golgi and secreted into the apoplast ^61^. The methyl groups are then selectively removed by pectin methylesterases (PMEs), which might be spatially regulated by pectic methylesterase inhibitors (PMEIs) ^60^. De-methylesterified HG chains are crosslinked into a tightly packed conformation by calcium bridges, which contribute to the wall strength ^32,62^. However, in some tissues decline in HG-calcium complexes has been correlated with a decrease in wall expansibility and an increase in wall stiffening^56,63^. Thus, the diversity and complexity in pectin structure and distribution in different tissues/cells and in different wall domains within a single cell might reflect functional diversity in the fine control of pectin gel rheology ^60^. Given the importance of pectin for PC morphogenesis (Fig. 3a, b)^7^, we further determined the pattern of pectin methylesterification in developing PCs (2-day cotyledons) by immunostaining with the LM20 and JIM5 monoclonal antibodies that recognize highly methylesterified and highly de-methylesterified pectin, respectively. The intensity of LM20 staining was greater in the lobe region when compared to the indentation area of the same cell, indicating a preferential distribution of highly methylesterified pectin in the lobe region (Fig. 3c, d). In contrast, higher levels of de-methylesterified pectin were found in the indentation region, comparing to the adjacent cell lobe region as indicated by the JIM5 immunofluorescence intensity (Fig. 3e, f). Note that this pectin staining pattern was observed in stage II PCs^20,64^ when PCs have just attained their jigsaw puzzle pattern and are still developing. PCs at early stages (undifferentiated and stage I cells) or late stage (fully expanded stage III) likely have very different cell wall structures and composition^20^. Indeed PCs on the late stage (the third leaves of 3-week old plants) did not exhibit a clear preference in the distribution of highly de-methylesterified and methylesterified pectin between the lobe and indentation regions (Fig. 3e, f). These results suggest that dynamic changes in pectin structure occur during the development of interdigitated PCs.

To investigate the functional significance of pectin de-methylesterification levels in PC morphogenesis, we characterized PC shape in mutants with reduced pectin de-methylesterification. We examined PCs phenotype in a knockout mutant for *PME3 (AT3G14310*), highly expressed in expanding cotyledons and true leaves^65 66^ and in a line overexpressing a pectin methylesterase inhibitor (PMEI1-OE)^67^. Both lines showed interdigitation defects similar to those of the *fer-4* mutant with simpler cell shapes and wider indentation necks (Fig. 3g, h). These results suggest that pectin methylesterifcation is spatially regulated at the subcellular level during the early stages of and is important for PC morphogenesis.

### FER preferentially binds de-methylesterified pectin

The significance of pectin methylesterifcation levels described above promoted us to determine its impact on pectin binding to FER. Interestingly FER proteins were readily pulled down by de-methylesterified pectin. Both full-length FER protein expressed from protoplasts (Fig. 3i) and FER-ECD recombinant protein produced from *E.coli* (Fig. 3j) interacted much more strongly with polygalacturonic acid (PGA, 0% methylesterification) compared with pectin extracted from apple (50-75% methylesterified) (Fig. 3i, j). Furthermore, *in vitro* pull-down assays showed that FER’s MALA domain preferentially bound de-methylesterifed pectin (PGA) compared to the highly methyesterified pectin (Supplementary Fig. 3a).

Since Ca^2+^ crosslinks de-methlyesterified HG via the carboxy groups, we investigated the importance of this conformation in binding with FER. We determined the binding of FER to PGA in the ionic buffer (0.5 mM CaCl_2_/150 mM NaCl) containing 5 mM, EDTA, a Ca^2+^ chelator, or in an ionic buffer, in which CaCl_2_ was replaced with 0.5 mM MgCl_2_. EDTA prevents the formation of PGA crosslinks, and Mg^2+^ ions are unable to stabilize PGA crosslinks due to their large size^62^. Both EDTA and MgCl_2_ treatments greatly suppressed the binding of FER to PGA (Supplementary Fig. 3b). Thus pectin crosslinks are critical for binding to FER. Taken together, our results suggest that FER favors binding de-methylesterified crosslinked pectin through the extracellular MALA domain to regulate PC morphogenesis.

### Pectin activates ROP6 signaling in a FER-dependent manner

Because both mutations in *FER* and reductions in pectin de-methylesterification result in a PC phenotype similar to that induced by ROP6 signaling defects^20^, we hypothesized that binding of de-methylesterified pectin to FER activates the ROP6 signaling pathway. We first determined the requirement of FER for ROP6 activation using an effector binding-based assay^68^. When the assay was conducted using an anti-ROP6 antibody, we observed dramatic ROP6 activity reduction in the *fer-4* mutant compared to wild-type (Fig. 4a, b). This reduction was further confirmed by another assay, in which GFP-ROP6 was introduced into Col-0 and the *fer-4* mutant and detected with an anti-GFP antibody (Supplementary Fig. 4a, b). Consistent with the changes in PC phenotypes of *pme3* and *PMEI1-OE* (Fig. 3g, h), ROP6 activation was also compromised in both *pme3* and *PMEI1-OE* lines (Fig. 4a, b). These results support the hypothesis that FER’s sensing of de-methylesterified pectin leads to the activation of the intracellular ROP6 signaling pathway. To further test this hypothesis, we determined whether pectin-mediated activation of ROP6 requires FER. Col-0 protoplasts were treated with PGA, and the activation of ROP6 was determined. Intriguingly, PGA treatment greatly induced ROP6 activation (Fig. 4c, d), and importantly this induction was compromised in *fer-4* mutant protoplasts (Fig. 4c, d). Hence PGA promotes ROP6 activation in a FER-dependent manner.

**Figure 4.**
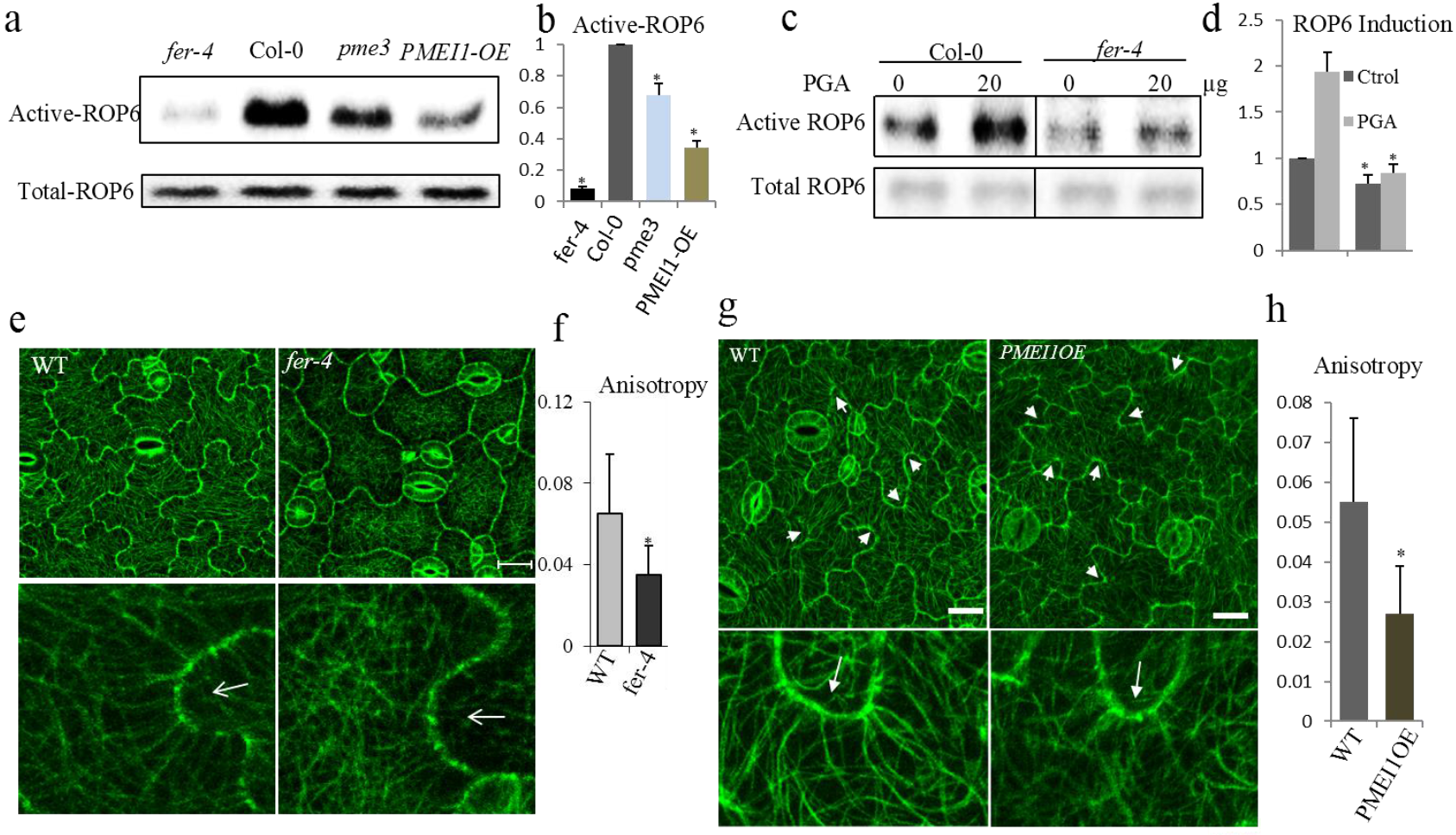
FER regulates PC morphogenesis through ROP signaling. **a and b**, Active ROP6 in wild-type, *fer-4, pme3* and *PMEI1-OE* mutants was analyzed by pull-down with the effector RIC1, as described previously (44), and the pulled down ROP6 was determined by a ROP6 antibody (a). The relative active of ROP6 level was quantified (b). 7 days seedlings were used in this assay. **c and d**, Activation of ROP6 by PGA in *Arabidopsis* protoplasts. ROP6 activities were determined in wild-type and *fer-4* protoplasts, isolated from 4 weeks adult plants, that were treated with or without 20 μg PGA (c). The relative active of ROP6 level was quantified (d). Data are mean activity levels from three independent experiments ±SE (b and d). **e and f**, PC cortical microtubules of wild-type (*GFP-MAP4*) and the *fer-4* mutant (*fer-4xGFP-MAP4*) (e). *Bottom* panel showed the magnified PC indentation regions. The degree of cortical microtubules anisotropy was quantified (f). Arrows indicate the indentation regions. Scale bar, 10 μ m. Data are mean degrees from 20 independent cells ±SE. **g and h**, PC cortical microtubules of wild-type (*GFP-TUB*) and the *PMEI1-OE* line (*PMEI1-OExGFP-TUB*) (e). *Bottom* panel showed the magnified PC indentation regions. The degree of cortical microtubules anisotropy was quantified (h). Arrows indicate the indentation regions. Scale bar, 50 μ m. Data are mean degrees from 20 independent cells ±SE. Asterisks indicate the significant difference (p≤0.05, Student’s *t*-test) between the wild type and the mutants in above assays.

The activation of ROP6 signaling promotes cortical microtubule organization to restrict lateral or radial cell expansion, thereby promoting the indentation of PCs^21^. In agreement with the FER-dependent activation of ROP6 by de-methylesterified pectin and the phenotypes of *fer-4* PCs with wider indentation necks (Fig. 1a, b and Supplementary Fig. 1c, d), *fer-4* PCs displayed less bundled and more randomly arranged cortical MTs, compared with wild-type cells (Fig. 4e, f). Furthermore, a similar reduction in the ordered arrangement of cortical MTs in the indentation region of PCs was found in the *PMEI1-OE* line (Fig. 4g, h, and Supplementary Fig. 4c, d) and *pme3* mutants (Supplementary Fig. 4c, d).

### RopGEF14 provides a direct link between FER and ROP6

ROPs are directly activated by ROP guanine nucleotide exchange factors (RopGEFs). RopGEF1 interacts with the FER kinase domain to activate ROP2 signaling in root hair development^39^. To assess which RopGEFs might be involved in PC morphogenesis, we analyzed the expression pattern of all 14 GEFs in different tissues, and found that only *RopGEFl, 6, 7*, and *14* transcripts were detected in cotyledons (Supplementary Fig. 5a). Furthermore, the characterization of PC morphogenesis of T-DNA insertion loss-of-function mutants for these RopGEFs revealed that two alleles of *gef14* exhibited a defect in PC interdigitation (Fig. 5 a, b, Supplementary Fig. 5b-i). Cotyledon PCs for these mutant alleles exhibited wider indentation compared to wild-type, reminiscent of a defect in the ROP6 signaling pathway^20^. The *gef14-2* PC indentation defect was completely rescued by complementing *gef14-2* with the *GEF14* cDNA fused with a Myc tag and its native promoter (Fig. 5a, b). Consistently, ROP6 activation (Fig. 5c, d, Supplementary Fig. 5j) and the ordering of cortical MTs were greatly compromised in the *gef14-2* mutant (Fig. 5e-g). Thus, we hypothesize that FER activates ROP6 signaling directly through RopGEF14 to regulate PC morphogenesis.

**Figure 5.**
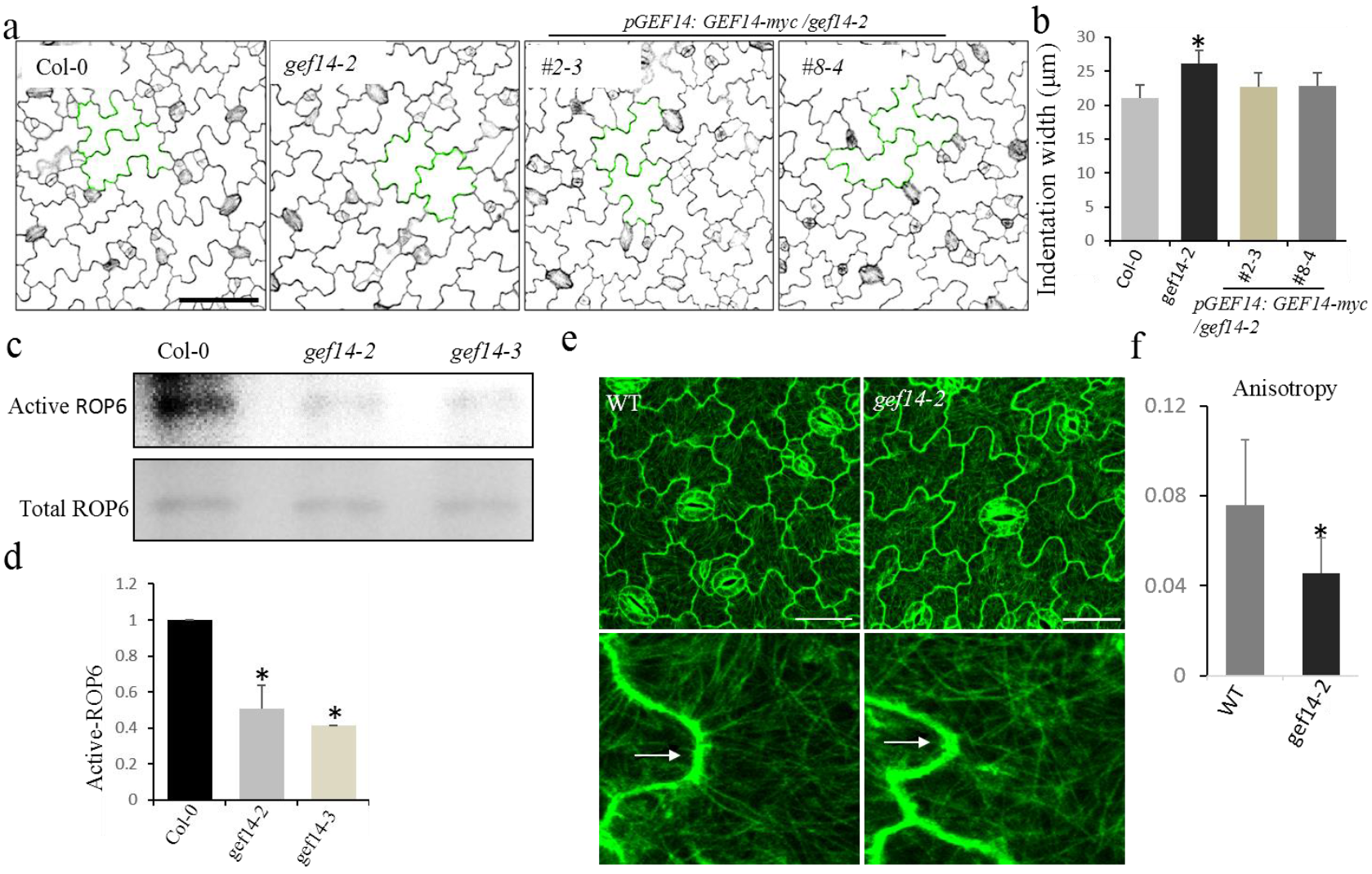
RopGEF14 regulates PC morphogenesis through ROP signaling. **a and b**, PC phenotypes of wild-type (Col-0), *gef14-2* and complementary line *pGEF::GEF-4xMyc/gef14-2* (a). The degree of PC interdigitation was determined by the width of indentation (b). Asterisk indicates a significant difference with p≤0.05 (Student’s *t*-test) between wild-type and the *gef14-2* mutant. Data are represented as mean ± SE. Scale bar, 50 μm. **c and d**, Activation of ROP6 in the wild-type and *gef14* mutants were analyzed by pull-down and determined by a ROP6 antibody (c). The relative active of ROP6 level was quantified (d). **e and f**, PC cortical microtubules of wild-type (*GFP-TUB*) and the *gef14-2* mutant (*GFP-TUB x gef14-2*) (e). *Bottom* panel showed the magnified PC indentation regions. The degree of cortical microtubules anisotropy was quantified (f). Arrows indicate the indentation regions. Scale bar, 50 μ m. Data are mean degrees from 20 independent cells ±SE. Asterisks indicate the significant difference (p≤0.05, Student’s *t*-test) between the wild type and the mutants in above assays.

To test this hypothesis, we first determined whether RopGEF14 forms a complex with FER and ROP6. We performed coimmunoprecipitation (CO-IP) assays in *pGEF14-GEF14-Myc* transgenic plants. As shown in Fig. 6 a and b, RopGFF14-Myc coimmunoprecipitated FER and ROP6, which were detected with α-FER^42^ and α-ROP6 antibodies, respectively (Fig. 6a, b). The association of FER with RopGEF14 was further confirmed using *35s-GFP-GEF14* transgenic plants. FER coimmunoprecipitated GFP-RopGFE14 but not GFP alone from transgenic plants (Supplementary Fig. 6a). These co-immunoprecipitation results are also consistent with those of yeast-two-hybrid assay suggesting a direct interaction between GEF14 and FER^39^. Moreover, recombinant MBP-GEF14 proteins specifically pulled down the full length of FER-HA and FER-TCD-HA expressed in mesophyll protoplasts (Supplementary Fig. 2b), but not FER-ECD-HA (Supplementary Fig. 2b), further confirming the interaction between RopGEF14 and FER’s intracellular kinase domain (Supplementary Fig. 6c).

We next conducted a series of experiments to confirm that ROP6 is also a part of the FER-RopGEF14 complex. Co-IP assays showed that ROP6 associated with RopGEF14 (Fig. 6b). MBP-GEF14 recombinant proteins efficiently pulled down ROP6-Flag expressed in protoplasts (Supplementary Fig 6d). Our co-IP assays further detected the association of FER with ROP6 in mesophyll protoplasts co-expressing FER-Flag and ROP-GFP (Fig. 6c). FER-Flag proteins were immunoprecipitated by ROP6-GFP using anti-GFP trap. Complementary results were obtained using FER-GFP to immunoprecipitate ROP6-Flag (Supplementary Fig. 6b). Taken together, these data indicate that FER regulates ROP6 signaling by directly interacting with the RopGEF14 /ROP6 complex.

Our genetic analysis further corroborates the physical interaction among FER, RopGEF14, and ROP6. Previously, we showed that a constitutively active ROP6 mutant (*CA-rop6*) dramatically suppressed lobe formation and increased the width of indentation necks in PCs^20^. PC phenotypes of the *CA-rop6 fer-4* double mutant was identical to those observed in *CA-rop6* (Supplementary Fig. 7a), suggesting that FER and ROP6 act in the same genetic pathway. Consistently with the reduction ROP6 signaling in the *fer-4* and *gef14-2* mutants (Fig. 4a-f, Fig. 5c-f), overexpression of ROP6 partially restored the *fer-4* and largely restored the *gef14-2* PC phenotypes, respectively (Fig. 6d-g). Furthermore, the PC indentation widths of *fer-5gef14-2* and *gef14-2rop6* double mutants were identical to those observed in *fer-5* and *gef14-2* mutants (Supplementary Fig. 7b-e). These results demonstrate that FER binds de-methylesterified pectin leading to the activation of the ROP6 signaling pathway through RopGEF14 and promotes the indentation during PC morphogenesis.

**Fig. 6.**
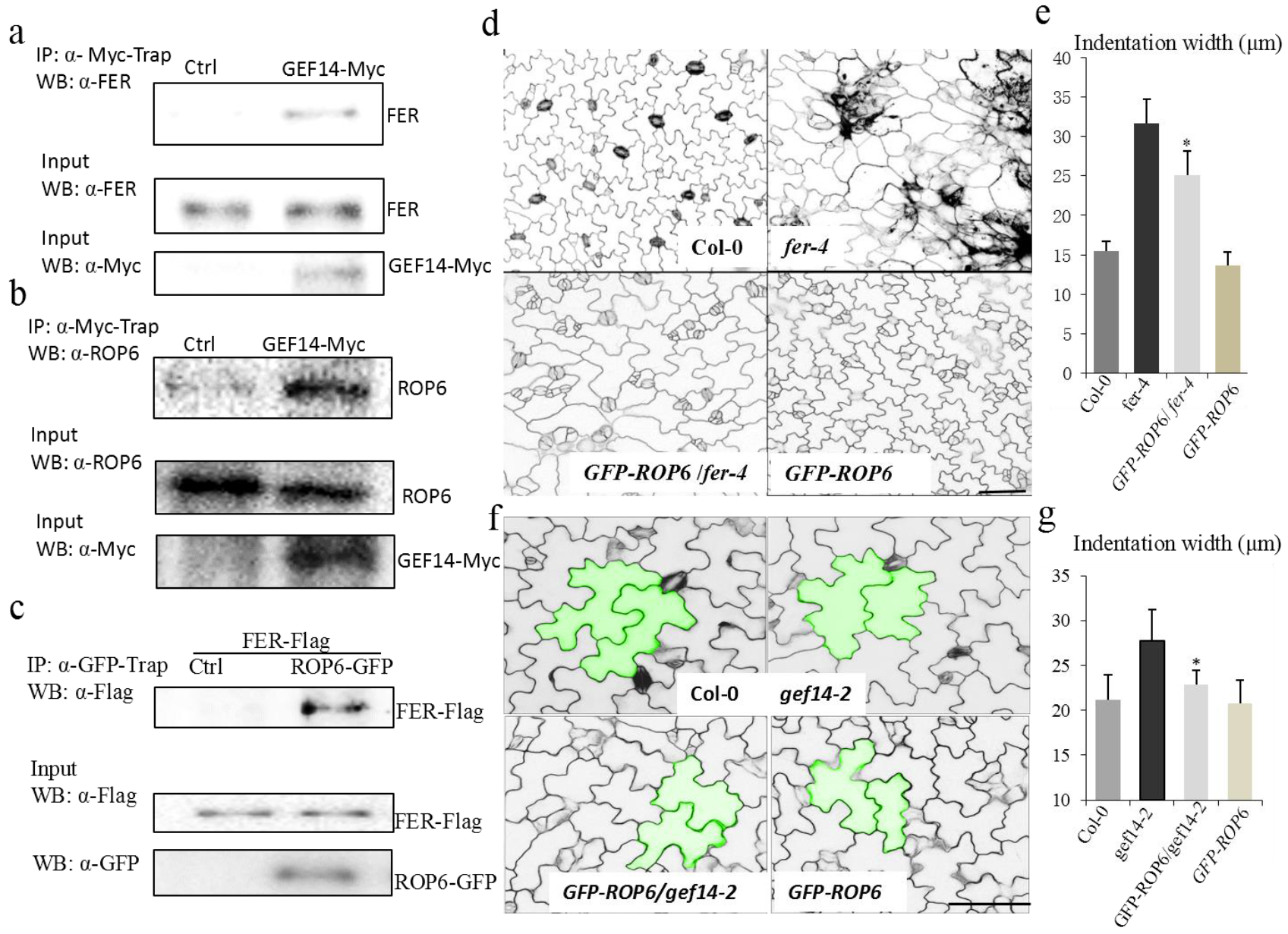
FER associates with RopGEF14 and ROP6 complex in regulating PC morphogenesis. **a**, FER associates with RopGEF14 in transgenic plants. Proteins from 10-day-old *pGEF14::GFE14-4xMyc/gef14-2* or *gef14-2* seedlings were immunoprecipitated with α-Myc-Trap antibody and analyzed with Western blot using α-FER antibody (Top). The expression of FER and GEF14-Myc in transgenic plants are shown (Middle and Bottom). **b**, ROP6 associates with RopGEF14 in transgenic plants. Proteins from 10-day-old *pGEF14::GFE14-4xMyc/gef14-2* or *gef14-2* seedlings were immunoprecipitated with α-Myc-Trap antibody and analyzed with Western blot using α-ROP6 antibody (Top). The expression of ROP6 and GEF14-Myc in transgenic plants are shown (Middle and Bottom). **c**, FER associates with ROP6 in protoplasts. Co-IP was carried out with an α-GFP-Trap antibody (IP: α-GFP-Trap), and the proteins were analyzed by using Western blot with α-Flag antibody. Top shows that FER-Flag coimmunoprecipitated with ROP6-GFP (IP: α-GFP-Trap; WB: α-Flag). Middle and Bottom show the expression of FER-Flag and ROP6-GFP proteins (WB: α-Flag or α-GFP for input control). **d and e**, ROP6 genetically acts downstream of FER in regulating PC morphogenesis. PC morphogenesis was characterized by 2 DAG wild-type, *fer-4, ROP6-GFP/fer-4 and ROP6-GFP* seedlings. Overexpression of ROP6-GFP partially restored the PCs defects of the *fer-4* mutant. Scale bar, 100 μ m. Asterisks indicate the significant difference (p≤0.05, Student’s *t*-test) between the *fer-4* and the *fer-4/ROP6-GFP* in above assays. **f and g**, ROP6 genetically acts downstream of RopGEF14 in regulating PC morphogenesis. PC morphogenesis was characterized by wild-type, *gef14-2, ROP6-GFP/gef14-2*, and *ROP6-GFP*. Overexpression of ROP6-GFP largely restored the PCs defects of the *gef14-2* mutant. Scale bar, 50 μ m. Asterisks indicate the significant difference (p≤0.05, Student’s *t*-test) between the *gef14-2* and the *ROP6-GFP/gef14-2* in above assays.

## DISCUSSION

Our comprehensive biochemical, genetic and cell biological data unequivocally demonstrate that upon sensing cell wall pectin via its extracellular domain the cell surface receptor kinase FER from the CrRLK1L subfamily directly activates ROP6 GTPase signaling and that this cell wall sensing/signaling pathway regulates PC shape formation in the *Arabidopsis* leaf epidermis. Therefore, FER is at least one of the long-sought cell wall sensors that couple a specific cell wall polysaccharide with intracellular signaling to regulate a particular cellular process. FER connects the cell wall properties to the control of cell expansion, likely by sensing dynamic changes in the cell wall composition and structure. Sensing the changes in pectin modification in the cell wall might be a common mechanism for the regulation of polar cell expansion. ANXUR1/2 and BUPS1/2 from the same family of FER control the cell wall integrity of elongating pollen tubes, which is also dependent on pectin de-methylesterification ^48,69^. FER-based monitoring of cell wall integrity and dynamic might also provide a mechanism to coordinate the cell growth between neighboring cells in plant tissues. In line with this, *fer* mutations disrupt coordinated cell growth causing random cell expansion in the root tip^70^ and severe defects in the interdigitated PC shape (Fig. 1a, b and Supplementary Fig. 1a, b). During interdigitated cell growth in PCs, the initial ROP2 activation might induce differential modifications of pectin between lobing and indenting sides (e.g, via secretion of PMEs, PMEIs, or pectinase), allowing the coordination between lobing and indenting sides via interactions between de-methylesterified pectin and FER.

By sensing specific wall components, FER and other CrRLK1L family members may directly monitor the cell wall integrity and activate compensatory pathways to maintain wall integrity. FER functions in plant defense response and is proposed to do so by monitoring the disruption of the cell wall caused by pathogen invasion^47^, though direct evidence for this role is lacking. Above mentioned THE1 is involved in sensing or signaling disturbances in cell wall cellulose biosynthesis to activate lignin biosynthesis^30^. Interestingly, yeast cells regulate cell wall integrity using a cell surface receptor-Rho GTPase signaling system^71" rid="c73">73^. The cell surface receptor-Rho GTPase signaling systems appear to be a common mechanism for surveying and regulating cell wall integrity.

By binding structural cell wall components such as de-methylesterified pectin, FER, and other CrRLK1L family members might sense the mechanical signals of either internal or external origin. This is consistent with a role for FER in mechanical signal transduction in *Arabidopsis*^70^. Finally, our findings expand the list of FER ligands including RALF^40,41^, suggesting that FER and likely its relatives may act as integrators of external signals. Clearly, CrRLK1L/RALF and LRX/RALF complexes are required for cell wall integrity maintenance in pollen tubes^7,48^ and roots^39^. Notably, the apical wall of pollen tubes is predominantly composed of pectin, with highly methylesterified pectin deposited at the growing tip and de-methylesterified pectin confined to the shank^69^. Reducing the activity of PME results in pollen tube rupture, reminiscent of the defect in the CrRLK1L/RALF and LRX/RALF pathways^69,74^. PME activity requires an alkaline pH optimum, while de-methylestrification of pectin leads to the cell wall acidification^75,76^. It is intriguing to propose that FER and other CrRLK1L members coordinate the sensing of RALFs (as alkalizing peptides) and pectin methylesterification levels to control the homeostasis of apoplastic pH and pectin status needed to maintain the cell wall integrity. How CrRLK1L members dance with RALFs and pectin to modulate cell growth, morphogenesis, and integrity maintenance is a fascinating question yet to be explored. Hence our findings here will propel a very fertile field of study on the sensing and signaling of cell wall dynamic, integrity and mechanics in plants.

## METHODS

### Plant Materials and Growth Conditions

The *fer-4* (GABI_GK106A06), *fer-5 (Salk_029056c*) were ordered from ABRC and *fer-2* was obtained from Ueli Grossniklaus (University of Zürich, Switzerland). The *pme3* and *PMEI1-OE* were obtained from Vincenzo Lionetti (Sapienza University of Rome, Italy) and the *arad1arad2* mutant was obtained from J. Paul Knok (University of Leeds, UK). The *GFP-MAP4xfer-4, GFP-TUBxPMEI1-OE, GFP-ROP6xfer-4, CA-rop6xfer-4, fer-5xgef14-2*, and *GFP-ROP6xgef14-2* mutants were generated by genetic crosses and confirmed by genotyping or Western blotting. *Arabidopsis* plants were grown in soil (Sungro S16-281) in a growth room at 23 °C, 40% relative humidity, and 75 μE m^−2.^s^−1^ light with a 12-h photoperiod for approximate 4 weeks before protoplast isolations. To grow *Arabidopsis* seedlings, the seeds were surface sterilized with 50% (vol/vol) bleach for 10 min, and then placed on the plates with 1/2 MS medium containing 0.5% sucrose, and 0.8% agar at pH 5.7. 2 to 6 days after germination (DAG) cotyledons were used for pavement cells characterization.

### Plasmid Construction and Generation of Transgenic Plants

Full-length and truncated variants *FER, BIR1*, and *ROP6* were amplified by PCR from Col-0 cDNA and cloned into a protoplast transient expression vector (obtained from Libo Shan & Ping He, Texas A&M) or plant expression vector pGWB641. The *FER* promoter of 1.3 kb was amplified by PCR from Col-0 genomic DNA and introduced into pGWB641 and pGWB605 binary vectors carrying FER-YFP and FER-GFP respectively. The truncated FER variants were constructed by overlapping PCR amplified from Col-0 cDNA and cloned into a protoplast compatible transient expression vector or plant expression vector pGWB642. FER-ECD and FER-MALA were amplified by PCR and cloned into proteins expression vector pDEST-HisMBP (obtained from *Addgene*). Full length of RopGEF14 was amplified from Col-0 genomic DNA and cloned into pGWB516. The GEF14 promoter of 1.5 kb together with GEF14 was amplified by PCR from Col-0 genomic DNA and introduced into pGWB516 binary vectors. Full length of RopGEF14 was amplified from Col-0 cDNA and cloned into pGWB506. All of the constructs were fully sequenced to verify mutations in the gene coding and promoter region. Stable transgenic lines were generated by using the standard *Agrobacterium tumefaciens*-mediated transformation in the *fer-4, gef14-2* mutants or Col-0.

### Confocal analysis of *Arabidopsis* cotyledons PC shape

*Arabidopsis* cotyledons were firstly incubated in PBS buffer with propidium iodide (PI) (2 mg/ml, 20 minutes). Then after washing with PBS buffer for three times, the samples were observed under confocal microscopy (Leica SP5 Laser Scanning Confocal.). The PCs images were taken at the mid-region of cotyledons. The widths of indentations were measured using *LAS AF Lite* software. Each dataset was generated from the measurement of at least 20 cells collected from 5 different cotyledons from 5 individual seedlings.

### GFP detection by proteins gel blotting and confocal laser scanning microscopy

Total proteins extraction and protein gel-blot analysis were performed as described before with modifications^28^. Soluble proteins were prepared using an extraction buffer (25 mM Tris-HCl, pH 7.5, 200 mM KCl, 5 mM MgCl_2_, and 5 mM DTT, 0.1% NP40) supplemented with a mixture of protease inhibitor (cOmplete Protease inhibitor cocktail; Roche). After washing and extraction for 15 times, the pellet was used for insoluble protein preparation. Insoluble proteins were extracted by boiling the pellet for 10 minutes in the extraction buffer (50 mM Tris-HCl, pH 6.8, 10% glycerol, 0.05% bromphenol blue and 50 mM DTT, 4% SDS). Total GFP was detected using a GFP polyclonal antibody (Santa Cruze, B-2, 1:1000 dilution) and subjected to horseradish peroxidase-conjugated mouse secondary antibody, and developed with enhanced chemiluminescence detection reagents (ThermoFish SuperSignal West Pico Chemiluminescent Substrate, Cat #. 34080). Ten-day-old seedlings were used for GFP detection. Whole cotyledons were directly mounted in PI (10 μ g/ml) or FM4-64 (5 μ g/ml) solution and observed with water objectives as described previously. GFP only transgenic seedlings were used as a control. GFP fluorescence and PI were excited simultaneously by a blue argon laser (10 mW, 488-nm blue excitation) and 535 nm for red excitation and detected at 515-530 nm for GFP and 600-617nm wavelengths for PI in a Leica SP5 Laser Scanning Confocal. For plasmolysis, cotyledons were incubated with PI for 20 minutes and washed with PBS buffer and subjected to a treatment of a 0.8 M Mannitol solution for 5 minutes before observation. Images were processed and arranged by ImageJ.

### Protoplast preparation and transient expression

Protoplasts were prepared according to the protocol described by Yoo et al^77^.Maxiprep DNA for transient expression was prepared using the Invitrogen PureLink Plasmid Maxiprep Kit. 2×10^5^ protoplasts were transfected with indicated FER-HA or truncated variants and incubated at room temperature for 10 hours. The protoplasts were collected and stored at −80 °C for further usage.

### Coimmunoprecipitate (Co-IP) assay

2 × 10^5^ protoplasts transfected with indicated plasmids were lysed with 0.5 mL of extraction buffer (10 mM Hepes at pH 7.5, 100 mM NaCl, 1 Mm EDTA, 10% (vol/vol) glycerol, 0.5% Triton X-100, and protease inhibitor mixture from Roche). After vortexing vigorously for 30 s, the samples were centrifuged at 12,470 × g for 10 min at 4 ° C. The supernatant was incubated with α-GFP-Trap antibody for 2 h with gentle shaking. The beads were collected and washed three times with washing buffer (10 mM Hepes at pH 7.5, 100 mM NaCl, 1 mM EDTA, 10% glycerol, and 0.1%Triton X-100) and once with 50 mM Tris • HCl at pH 7.5. The immunoprecipitated proteins were analyzed by Western blot with α-GFP or α-FLAG antibody. For seedling Co-IP, approximate 1 g of 10-day seedlings were ground in liquid N2 and further ground in 0.5 mL of ice-cold Co-IP buffer. Samples were centrifuged at 12,470 × g for 10 min at 4 °C. The resulting supernatant was used to perform the Co-IP assay with the same procedures as protoplast Co-IP assay with α-Myc-Trap or α-FER (Generated and purified from Rabbit)^42^ antibodies.

### *In vitro* pull-down assay

2 × 10^5^ protoplasts were lysed with 1 mL extraction buffer (20 mM Tris pH 8.2, NaCl 150 mM, 0.5% Triton X-100, 0.5 mM CaCl_2_ and protease inhibitor mixture from Roche)^78^. After vortexing vigorously for 30 seconds, the samples were centrifuged at 12000 × g for 10 minutes at 4 °C. 20 μl of supernatant was used as input control and the remainder of the supernatant was incubated with the indicated amounts of pectin or PGA for 2 −4 hours at 4 °C with gentle shaking. Pectin or PGA were pelleted and collected by centrifuging at 6000 x g for 10 minutes at 4 °C and washed three times with washing buffer (20 mM Tris pH 8.2, NaCl 150 mM, 0.5 mM CaCl_2_ and protease inhibitor mixture from Roche). 20 μl 1× SDS loading buffer was added to the pull-down pellet and the released proteins were analyzed by Western blot with a α-HA antibody.

Expression of MBP fusion proteins and affinity purification were performed as standard protocol. The protein concentration was determined with NanoDrop ND-1000 spectrophotometer and confirmed by the Bio-Rad Quick Start Bradford Dye Reagent. 200 ng *E.coli* (Rosetta DE3) produced recombinant proteins were incubated with 200 μl extraction buffer for 30 minutes and then centrifuged at 6000 × g for 10 minutes at 4 °C, 20 μl of supernatant was used as input control and the remainder of the supernatant was incubated with the indicated amounts of pectin or PGA for 2 hours at room temperature with gentle shaking. Pull-down was carried out as described above, and the pull-down proteins were determined by western blot with α-His or α-MBP antibodies.

### Immunolocalization of pectin and MT in PCs

Two DAG cotyledons or the epidermal layer from 3 weeks old plants the third pair leaves were fixed, frozen, shattered and permeabilized as described by Burn, J.E^34^. For pectin immunolocalization, samples were incubated with the primary antibodies JIM5 (1: 100, PlantProbes, University of Leeds, UK) and LM20 (1: 100, PlantProbes, University of Leeds, UK). Samples were then transferred to a buffer with the secondary antibody (FITC-conjugated anti-Rat IgG at 1: 100 dilution, Sigma). For MT immunolocalization, following incubation with a monoclonal anti-α-tubulin-FITC antibody produced in mouse (Sigma) and washed three times after incubation, the samples were observed under Leica SP5 Laser Scanning Confocal. For quantification of pectin in lobes and indentations, the fluorescence intensity was analyzed by ImageJ. The mean ratio of indention/lobe was calculated to indicate the differential distribution of de-esterified pectin and esterified pectin in lobe and indentation regions of PC.

### Quantitative analysis of cortical microtubule orientation

The *fer-4* mutant was crossed to the *GFP-MAP4* line and the *PMEI1-OE* was crossed to *GFP-TUB* line to enable the visualization of MTs. Images of PC MTs were generated using a Leica SP5 Laser Scanning Confocal. FibrilTool, which is an ImageJ plug-in to quantify fibrillar structures, was used to analyze the average anisotropy of MT^79^. The indentation regions were selected with the Polygon tool. With regard to the anisotropy score, “0” indicates no order (purely isotropic arrays) and “1” indicates perfectly ordered (purely anisotropic arrays). Each data set was from the measurement of at least 20 cells collected from 3 different cotyledons from 3 individual seedlings.

### Measurement of ROP6 activity

ROP activity measurement was performed as described by Xu^68^. 10-day-old 1/2 MS grown seedlings or protoplasts isolated from 4-week-old plants were used in this assay. Total ROP6 and activated ROP6 proteins that pull down by MBP-RIC1 (GTP-bound ROP6) were detected by Western blots using ROP6 or GFP and horseradish peroxidase-conjugated rabbit antibodies. The proteins levels were determined by ImageJ.

### RT-PCR Analysis

Total RNA was isolated from wild-type or mutant leaves or seedlings with TRIzol Reagent (Invitrogen). First-strand cDNA was synthesized from 1 μ g of total RNA with reverse transcriptase. The RT-PCR analysis was carried out by using the synthesized cDNA as templates, and Actin 8 was used as a control gene. The RT-PCR primer sequences are listed below.

### Primers for construct cloning and genotyping

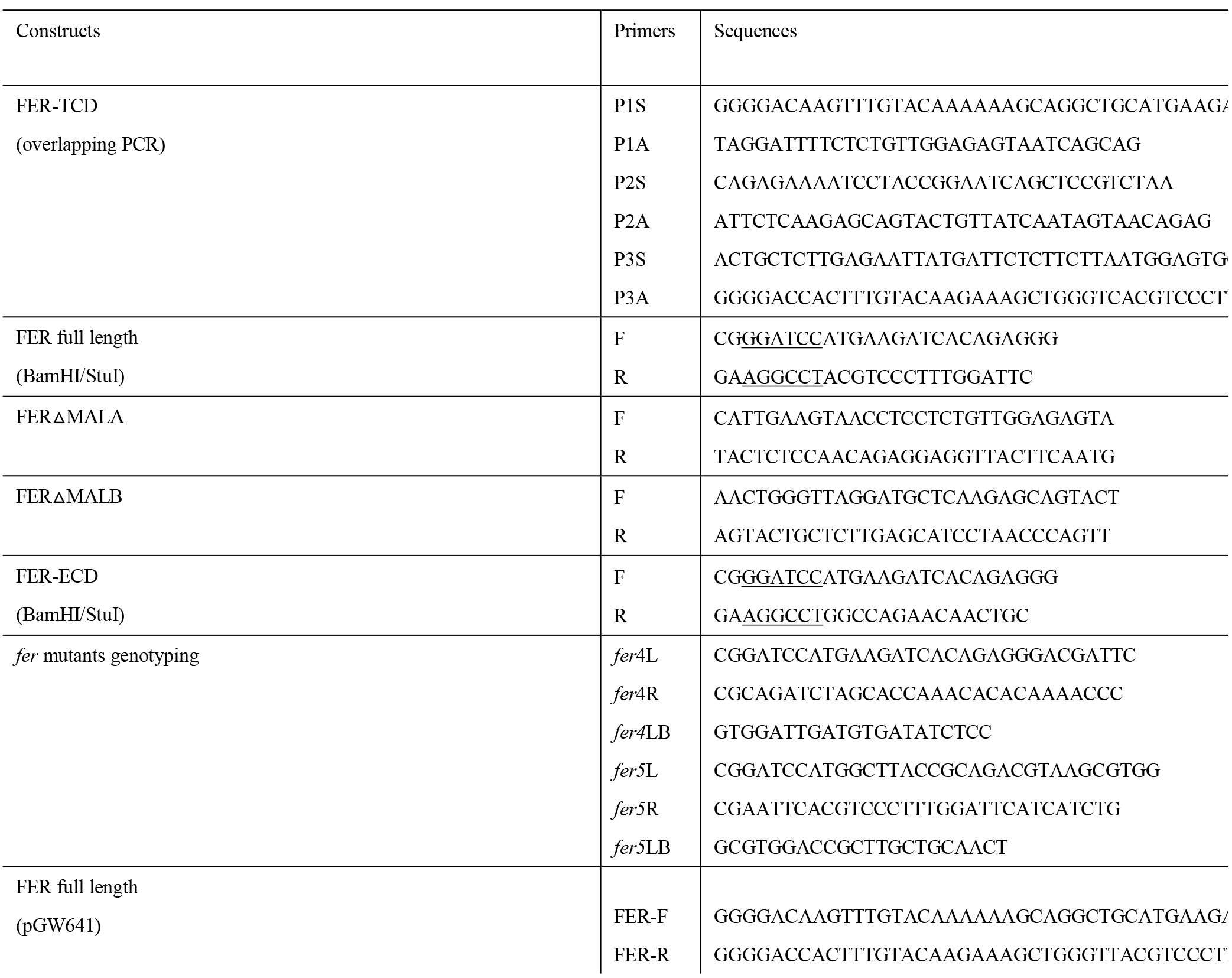

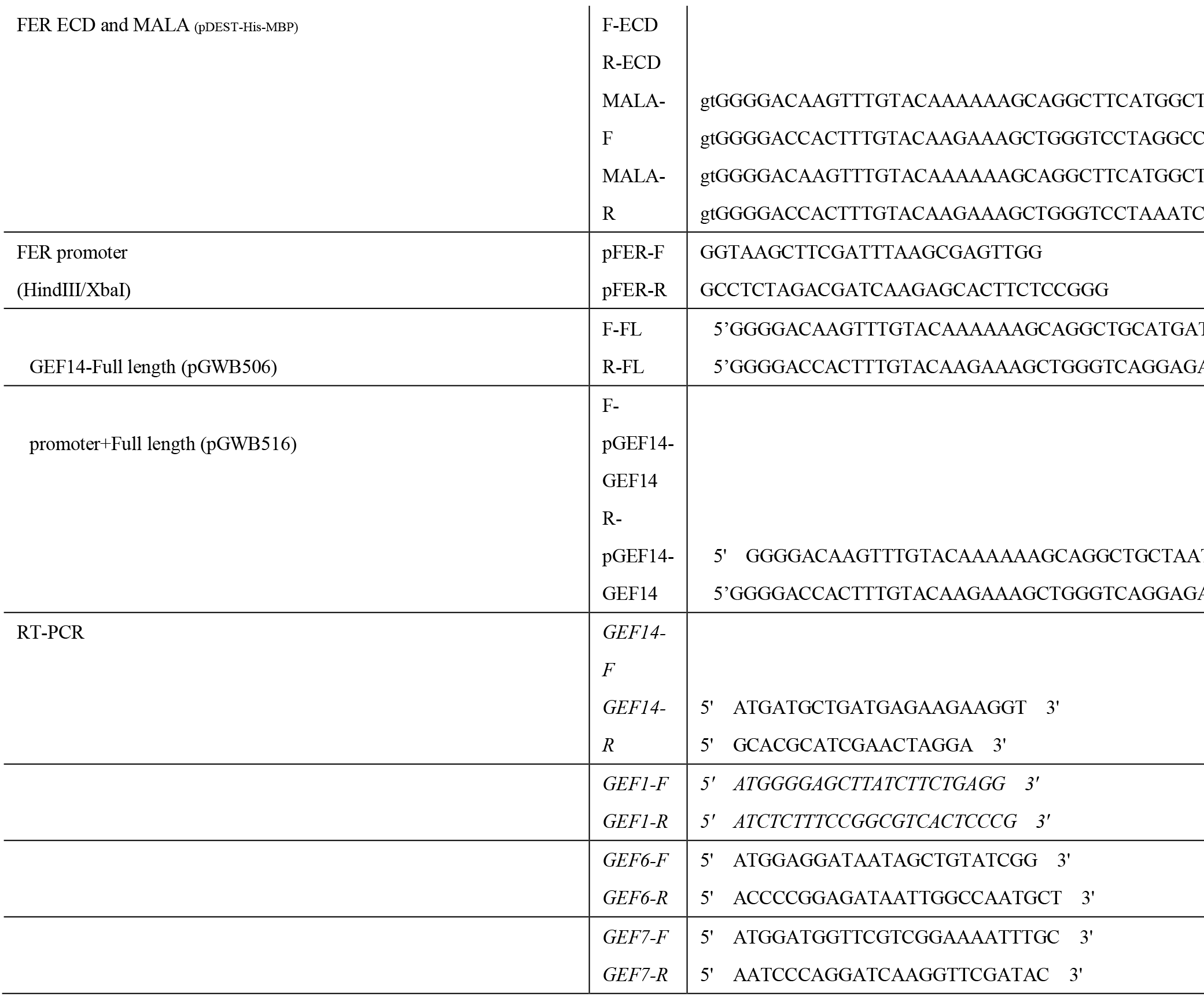

## ACKNOWLEDGEMENTS

We are grateful to Ueli Grossniklaus (University of Zürich, Switzerland) for *fer-2* seeds, to Vincenzo Lionetti (Sapienza University of Rome, Italy) for *pme3* and PMEI1-OE seeds and to J. Paul Knok (University of Leeds, UK) for *arad1 arad2* seeds. We gratefully acknowledge Eugene Nothnagel (University of California, Riverside, USA) for helpful discussions. We thank members of the Yang laboratory for their technical assistance and helpful discussions. This work is supported by a grant from the U.S. National Institute of General Medical Sciences to Z.Y. (GM081451) and by Fujian Agriculture and Forestry University. C. T. A. is supported by the Center for Lignocellulose Structure and Formation, an Energy Frontier Research Center funded by the U.S. Department of Energy, Office of Science, Basic Energy Sciences (award no. DE-SC0001090).

## Supplemental Data

**Supplemental Figure l.**
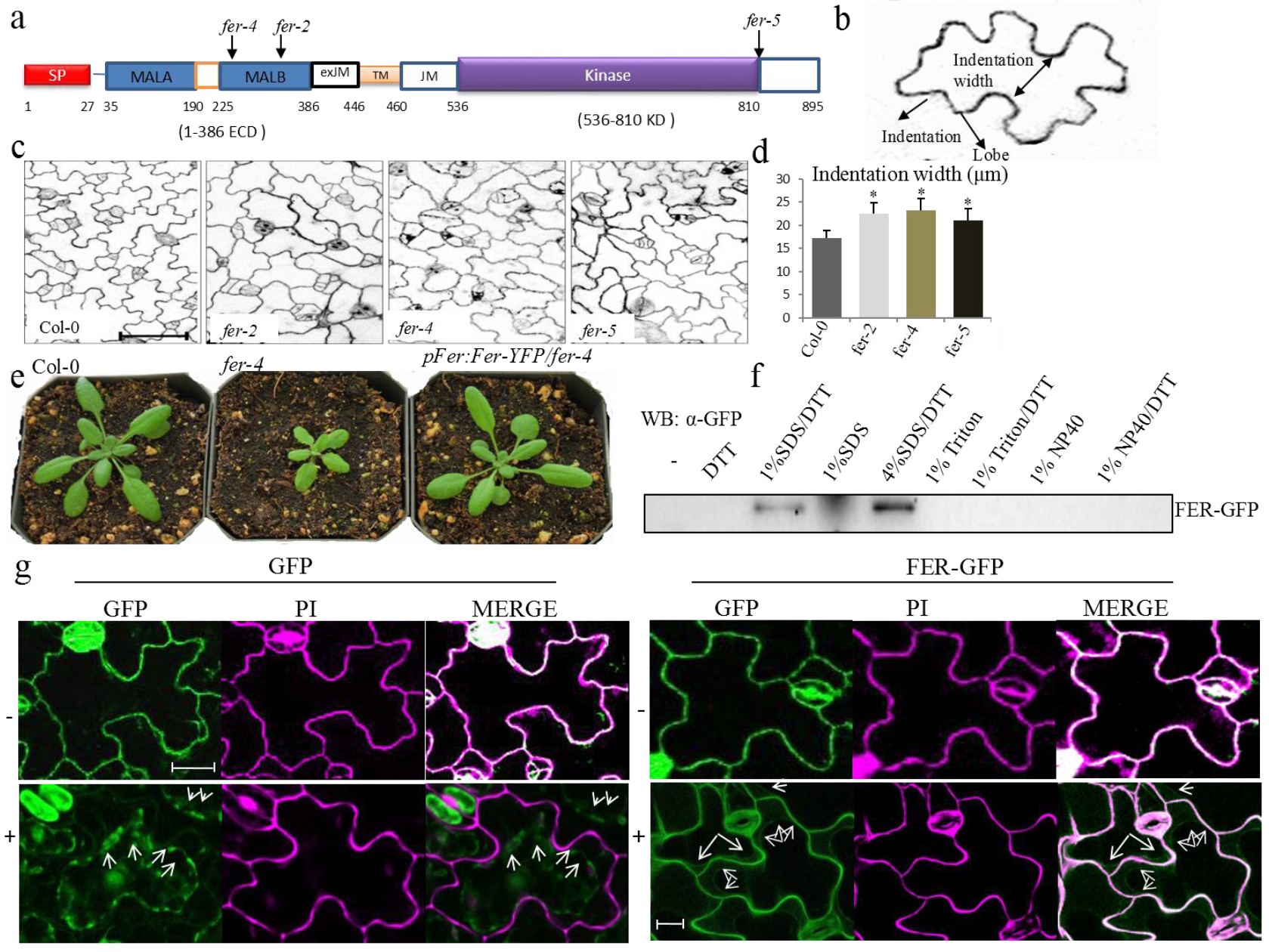
FERONIA is a cell wall-associated protein required for PC morphogenesis. **a**, Diagram of FER’s predicted domains and T-DNA insertion sites of 3 alleles of *fer* mutants (*fer-2, fer-4, fer-5*). **b**, A diagram depicting how the neck widths were measured. A dashed line segment was drawn between two indentations region, and the distance was decided as the indention width. **c**, Pavement cell phenotypes of the wild-type and *fer-2, fer-4* and *fer-5* mutants. Scale bars, 100 μm. 2 DAG seedlings cotyledon pavement cells were characterized. **d**, The degree of pavement cell interdigitation was quantified by determining the average indentation widths (Indentation width/ μm). The stars indicate the average indentation widths were significantly different (p≤0.05) between wild-type and the *fer* mutants. All data are represented as mean ± SE. **e**, The morphologies of 4 weeks old soil-grown wild-type, fer-4, and fer-4 complementation line plants. **f**, FER can be extracted by 1% SDS, 50 mM DTT from the cell fraction. The FER-GFP proteins were determined by Western blot with a GFP antibody. **g**, The cell wall localization of FER-GFP. (Top panels) GFP fluorescence was along with the cell surface of PCs before plasmolysis (−) indicated by overlapping signal with PI (Merge). During plasmolysis (+) GFP fluorescence localized with the cytoplasm (Bottom left panel). A small portion of the FER-GFP signal retreated with the cytoplasm and majority of the signal remained on the cell surface (Bottom right panel). Arrows indicate the regressed plasma membrane. Scale bar, 5 μm

**Supplemental Figure 2.**
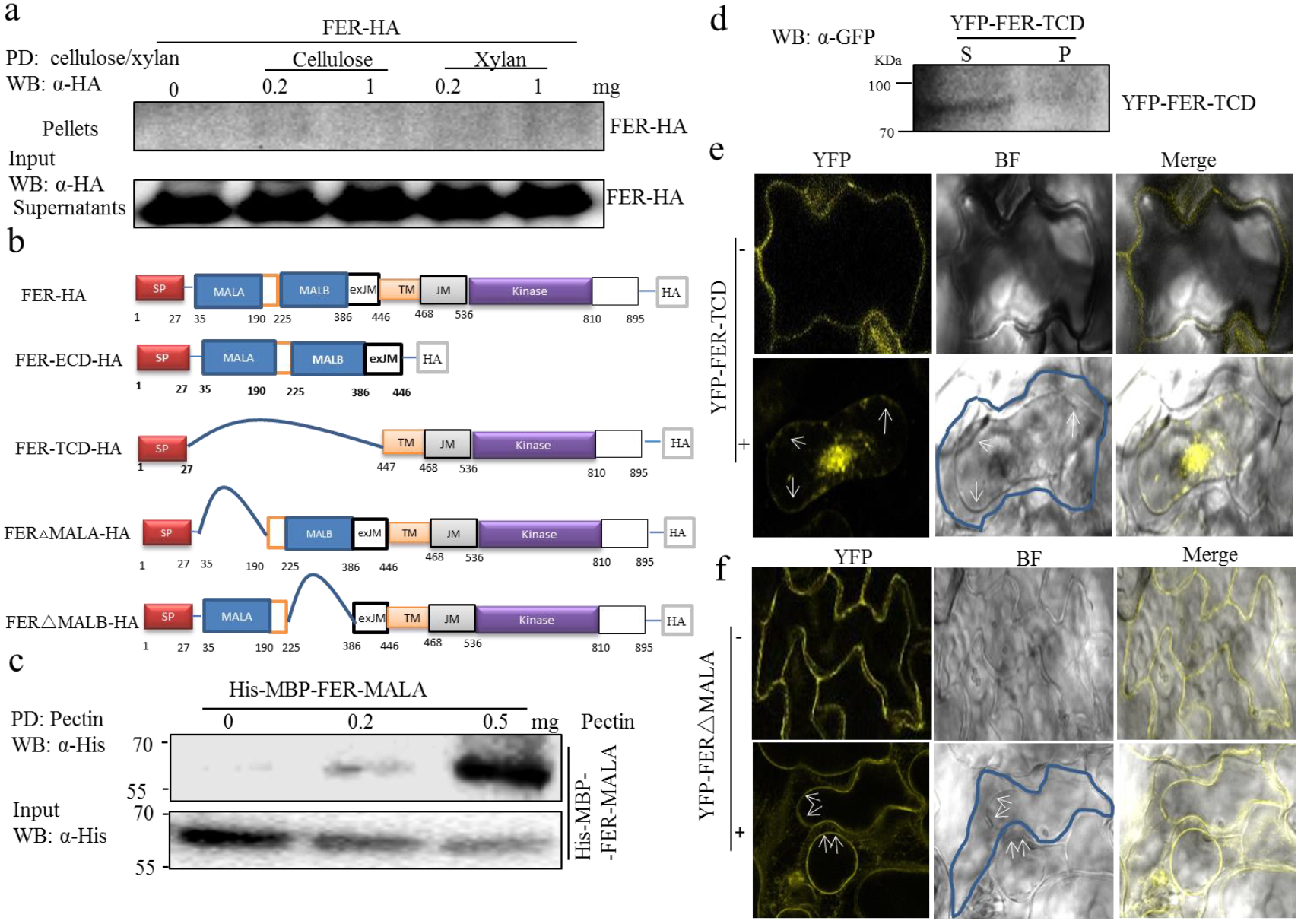
FER associates with pectin and cell wall with the MALA domain. **a**, FER failed to be pulled down by cellulose and xylan *in vitro*. FER-HA was expressed in *Arabidopsis* protoplasts. Pull-down (PD) was carried out with cellulose and xylan with indicated concentration, respectively. The FER-HA proteins were determined by Western blot with α-HA antibody. *Top* shows that FER-HA was not pulled-down by cellulose or xylan (PD: cellulose/xylan; WB: α-HA). *Bottom* shows the expression of FER-HA proteins (WB: α-HA as input control). **b**, Diagram of the full length and specific truncated FER-HA constructs. The FER protein contains a signal peptide domain (SP), a malectin-like extracellular domain A (MALA), a malectin-like extracellular domain B (MALB), an extracellular juxta-membrane domain (exJM), a transmembrane domain (TM), a juxta-membrane domain (JM) and a cytosolic kinase domain (Kinase). Numbers are the specific amino acid delineations between domains. FER-ECD-HA, construct that with the extracellular domain (ECD, includes SP, MALA, MALB, and exJM) of FER fuses with HA tag, the transmembrane domain and cytosolic domain (JM and Kinase domain) were deleted. The FER-TCD-HA construct contains FER signal peptide, a transmembrane domain and cytosolic domain (TCD, includes SP, TM, JM, and Kinase domain), with the extracellular domain (ECD) deleted and fused with HA tag at the C-terminus. The FER△MALA-HA construct has the FER MALA domain deleted and an HA tag at the C terminal. The FER△MALB-HA construct has the FER MALB domain deleted and an HA tag at the C-terminus. **c**, MALA domain directly binds to pectin *in vitro*. His-MBP-FER-MALA was purified from *E.coli*. PD was carried out as above and the proteins were determined by Western blot with α-His antibody. His-MBP-FER-MALA pull-down by pectin (Top). The input His-MBP-FER-ECD proteins (*Bottom*). **d**, Identification of YFP-FER-TCD proteins by Western blot within different fractions. Total soluble (S) and insoluble (P) proteins were prepared from seedlings of *35S::YFP-FER-TCD* transgenic plants and analyzed with α-GFP antibody by Western blot. **e**, Deletion of extracellular domain results in loss of cell wall binding of FER. Determination of the subcellular localization of YFP-FER-TCD by plasmolysis assay with (+) or without (−) a 0.8 M mannitol solution. Arrows indicated the plasma membrane region. **f**, MALA domain is required for FER cell wall binding. Determination of the subcellular localization of YFP-FER△MALA by plasmolysis assay with (+) or without (−) a 0.8 M mannitol solution as was described before. Arrows indicated the plasma membrane region.

**Supplemental Figure 3.**
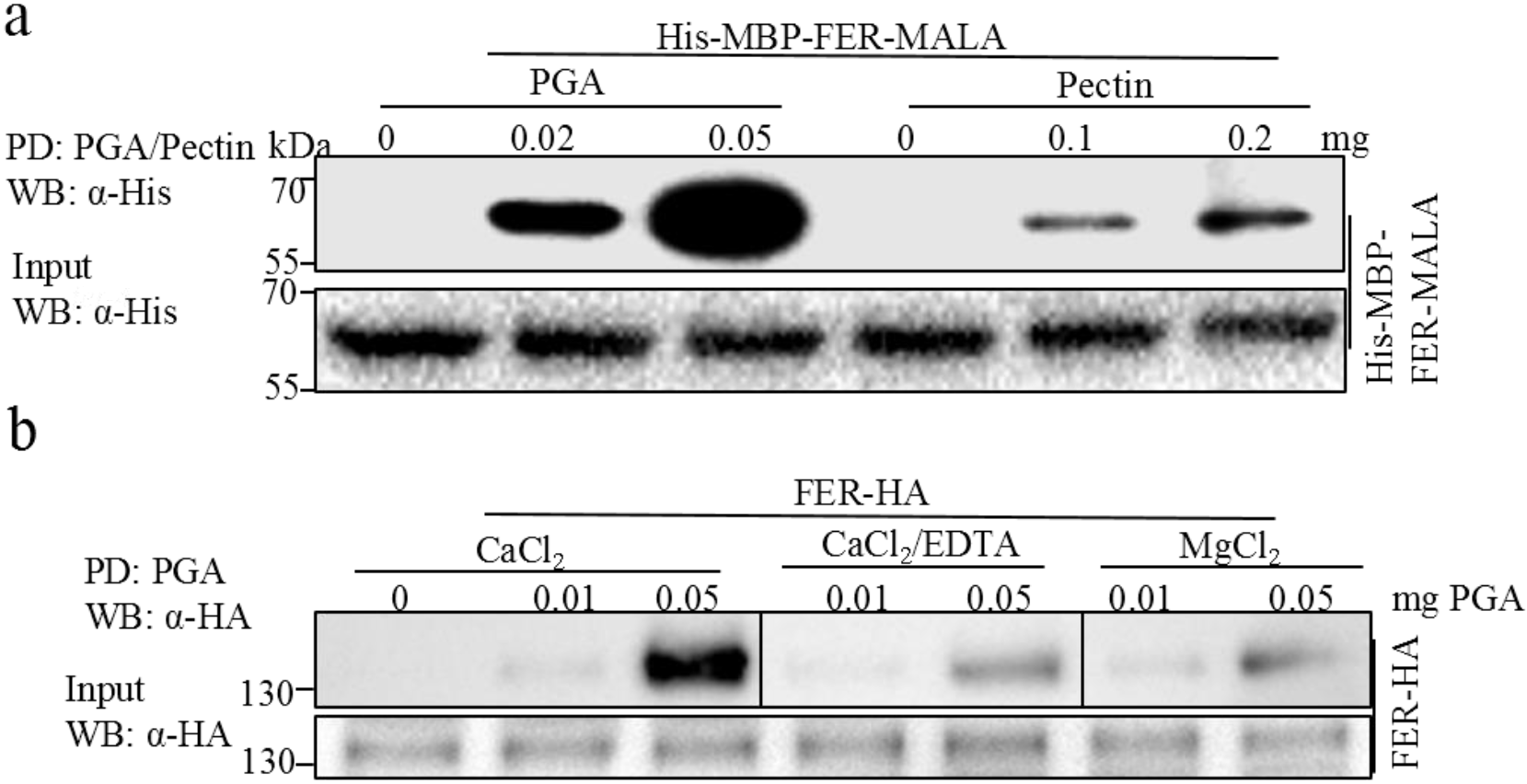
FER preferentially binds to PGA than pectin. **a**, FER MALA domain was associated with PGA and pectin *in vitro*. Pull-down was carried out with PGA and pectin as described previously. (*Top*) His-MBP-FER-MALA proteins produced from *E.coli* were pull-down by PGA or pectin. (*Bottom*) The His-MBP-FER-MALA proteins were determined by WB as input control. **b**, HG crosslink formation is required for FER-PGA association. An interaction between FER-HA and PGA was determined in different ionic solutions of concentrations as indicated. The pull-down was performed as described previously. *Top* panel shows that FER-HA pull-down by PGA (PD: PGA; WB: α-HA). *Bottom* panel shows the expression of FER-HA proteins (WB: α-HA for input control).

**Supplemental Figure 4.**
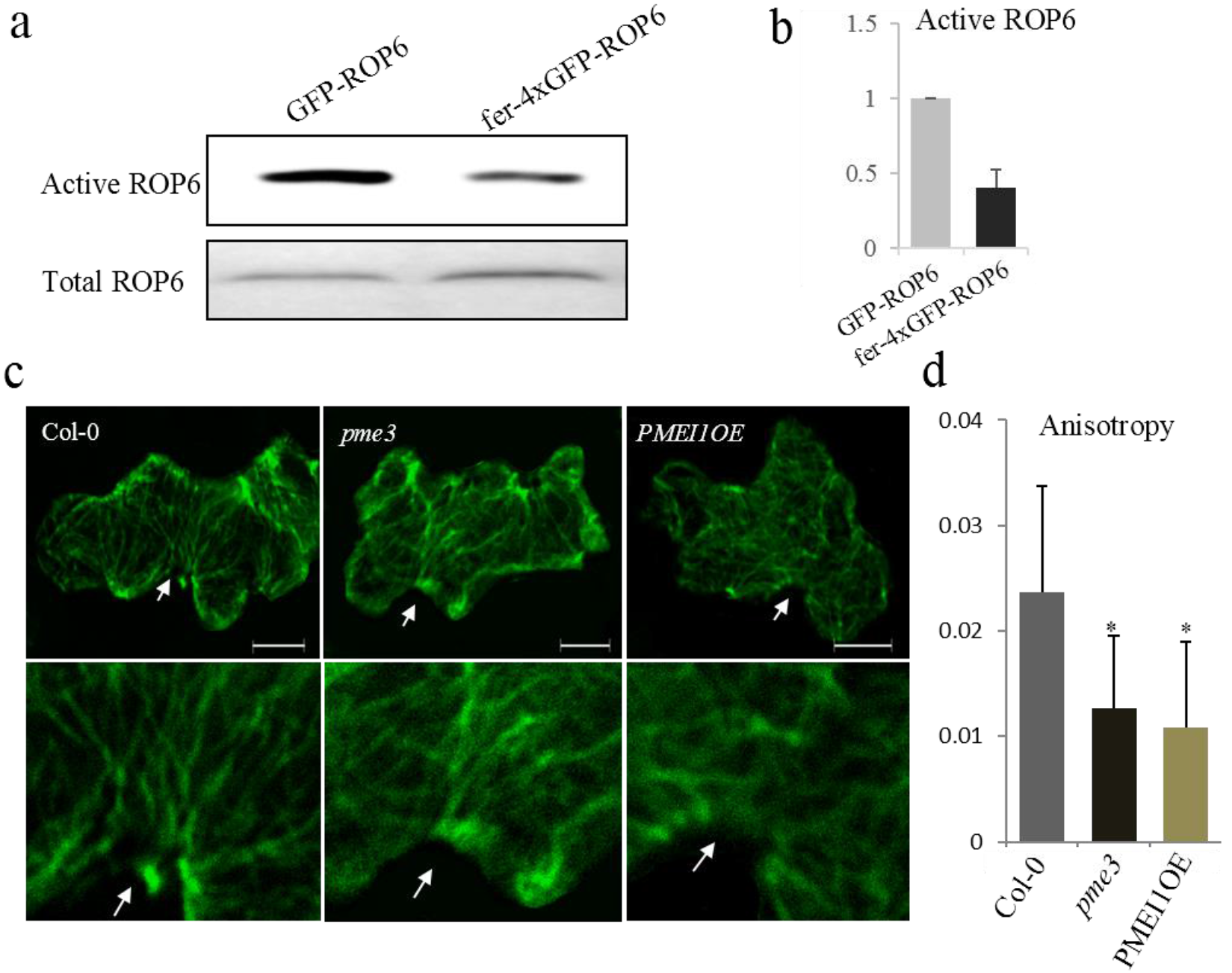
FER mediates PCs morphogenesis via ROP6 signal pathway. **a and b**, Activation of ROP6 in the wild-type (*GFP-ROP6*) and the *fer-4* mutant (*fer-4xGFP-ROP6*) were analyzed by pull-down assay and determined by GFP antibody (a). The relative active of ROP6 level was quantified (b). Protoplasts isolated from 4 weeks old adult plants were used in this assay. **c**, Cortical microtubule orientation defects of *pme3* and *PMEI1-OE* mutants. Wild-type PCs show highly ordered transverse cortical microtubules (with high anisotropy) in the indentation region, while microtubule arrays in the *pme3* and *PMEI1-OE* mutant were mostly present random orientations (Top panel). Magnified the PC indentation regions (Bottom panel). The microtubules were visualized by immunostaining with an anti-tubulin antibody. Arrows indicate the indentation region. Scale bar, 10 μm. **d**, The histogram shows that the degree of cortical microtubule anisotropy in wild-type and the *pme3* and *PMEI1-OE* mutants. Anisotropy was calculated using ImageJ. Data are mean degrees from 10 independent cells ±SE. Asterisks indicate the significant difference (p≤0.05, Student’s *t*-test) between the wild type and the mutants in above assays.

**Supplemental Figure 5.**
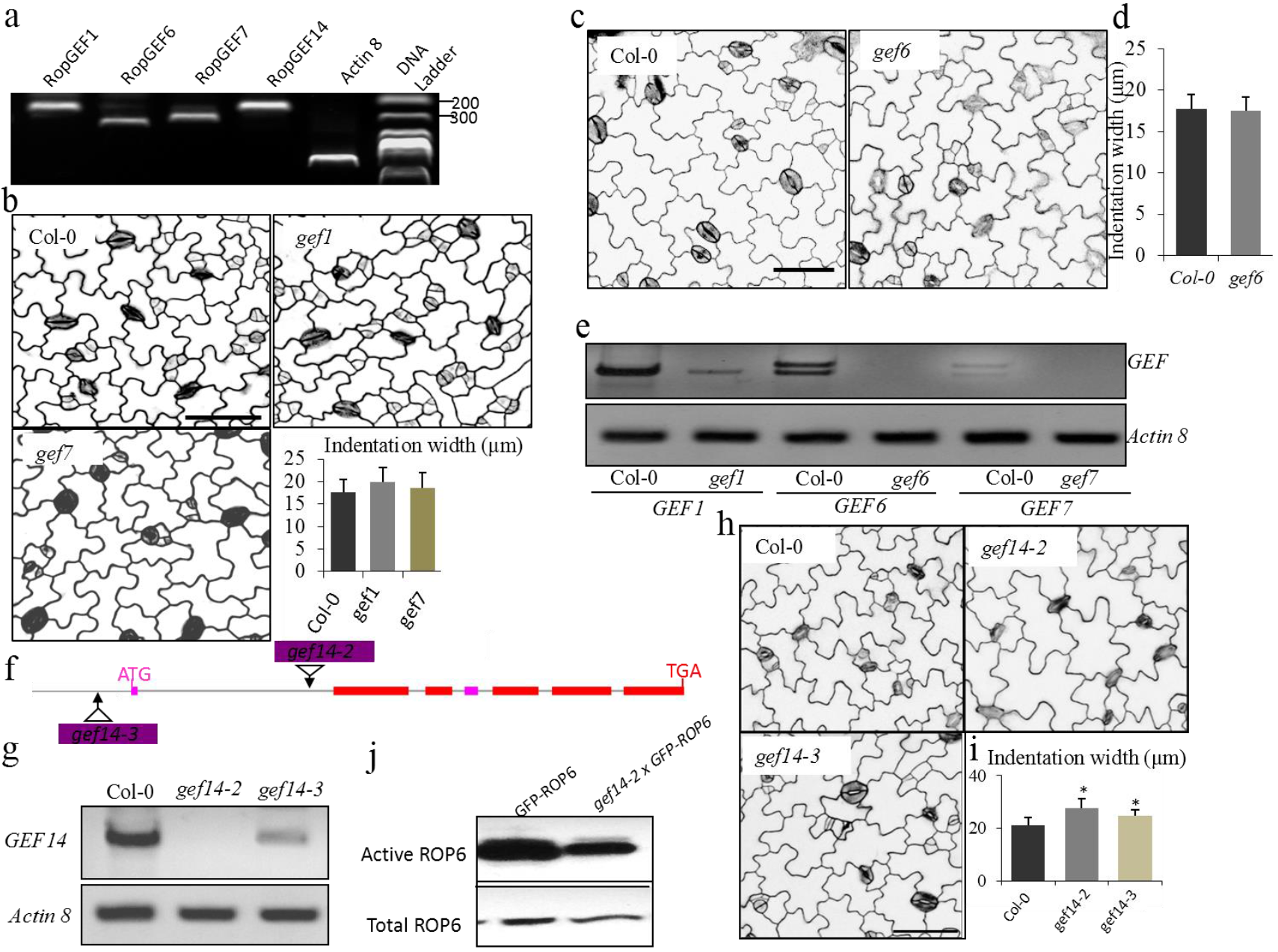
RopGEF14 is required for ROP GTPase activation in regulating PC morphogenesis. **a**, The expressions of *RopGEF1, 6, 7* and *14* in cotyledon were determined by RT-PCR. Actin 8 was included as an expression control. **b**, PC phenotypes of wild types (Col-0), the *gef1* and *gef7* mutants. The degree of 2 DAG PC interdigitation was determined by the width of indentation. **c and d**, PC phenotypes of wild types (Col-0), and the *gef6* mutants (c). The degree of 2 DAG PC interdigitation was determined by the width of indentation (d). **e**, RT-PCR analysis of *GEF1, GEF6, GEF7* and *Actin8* (control) in wild-type (Col-0) and the *gef1, gef6* and *gef7* T-DNA insertion mutants. **f**, T-DNA insertion sites in the *gef14* mutants with exons (red boxes). **g**, RT-PCR analysis of *GEF14*, and Actin8 (control) in wild-type (Col-0) the *gef14-2* and *gef14-3* T-DNA insertion mutants. **h and i**, PC phenotypes of wild types (Col-0), the *gef14-2* and *gef14-3* mutants (g). The degree of 3 DAG PC interdigitation was determined by the width of indentation (h). **j**, Activation of ROP6 in the wild-type (*GFP-ROP6*) and *gef14-2 (gef14-2xGFP-ROP6*) mutant were analyzed by pull-down and determined by a GFP antibody. Protoplasts isolated from 4 weeks old adult plants were used in this assay.

**Supplemental Figure 6.**
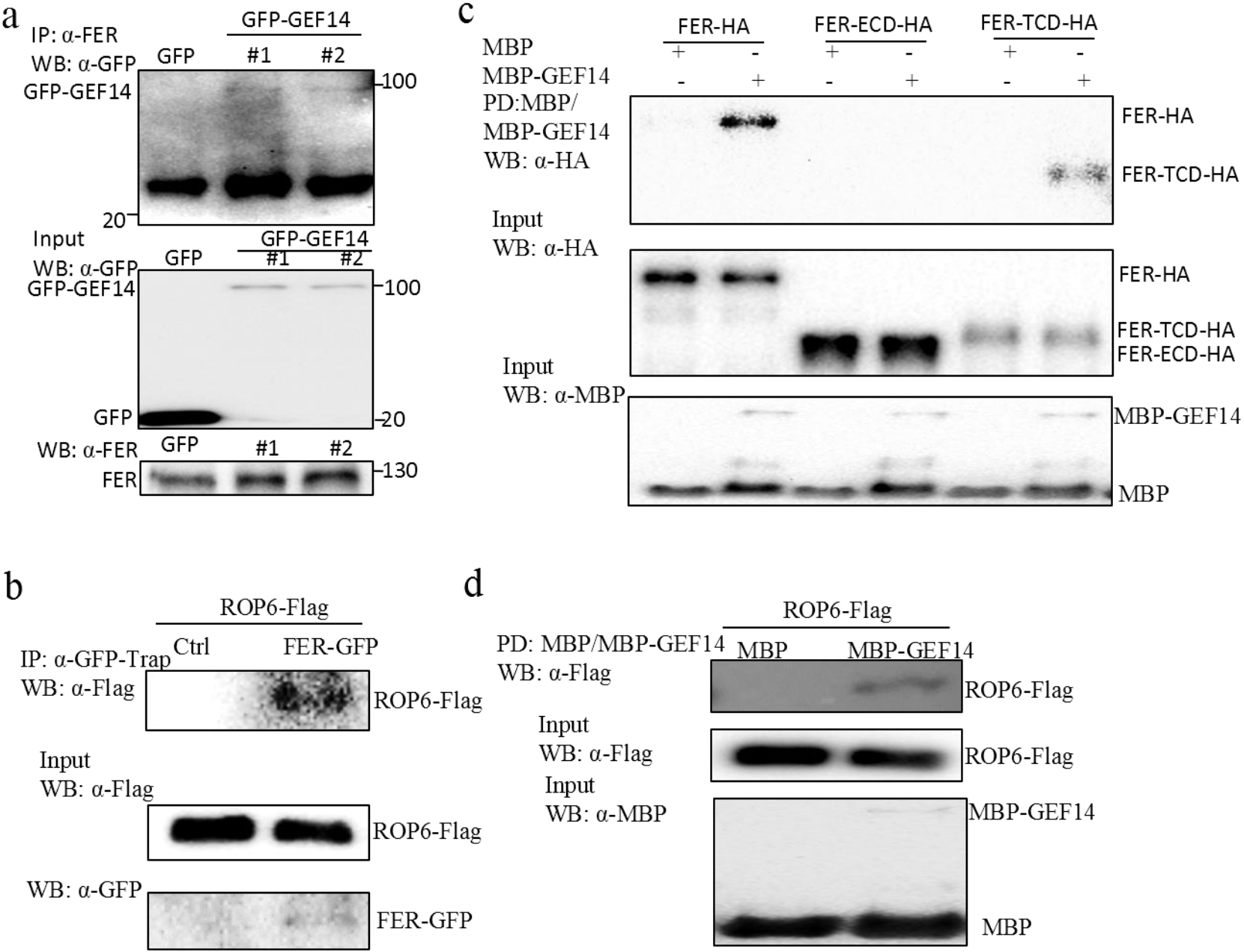
FER, GEF14, and ROP6 genetically act in the same signaling pathway in regulating PC morphogenesis. **a**, FER associates with RopGEF14 in transgenic plants. Proteins from 10-day-old *35S::GFP-GFE14* or *35S::GFP* transgenic seedlings were immunoprecipitated with α-FER antibody and analyzed with Western blot using α-GFP antibody (Top). The expression of GFP-GEF14, GFP, and FER in transgenic plants are shown (Middle and Bottom). **b**, FER associates with ROP6 in protoplasts. Co-IP was carried out with a α-GFP-Trap antibody (IP: α-GFP-Trap), and the proteins were analyzed by using Western blot with the α-Flag antibody. Top shows that ROP6-Flag coimmunoprecipitated with FER-GFP (IP: α-GFP-Trap; WB: α-Flag). Middle and Bottom show the expression of ROP6-Flag and FER-GFP proteins (WB: α-Flag or α-GFP for input control). **c**, FER associates with RopGEF14 through intracellular kinase domain. The full length of FER-HA, truncated FER-ECD-HA and FER-TCD-HA were expressed in protoplasts and pull down was carried out with recombinant MBP or MBP-GEF14 proteins (PD: α-MBP/MBP-GEF14), and the proteins were analyzed by using Western blot with α-HA antibody. Top shows that FER-HA and FER-TCD-HA coimmunoprecipitated with MBP-GEF14 (PD: α-MBP/MBP-GEF14; WB: α-HA). Middle shows the expression of FER-HA, FER-ECD-HA and FER-TCD-HA proteins (WB: α-HA for input control), and Bottom showed the input recombinant proteins (WB: α-MBP, for input control). **d**, ROP6 associates with RopGEF14. ROP6-Flag was expressed in protoplasts and pull down was carried out with recombinant MBP or MBP-GEF14 proteins (PD: α-MBP/MBP-GEF14), and the proteins were analyzed by using Western blot with α-Flag antibody. Top shows that ROP6-Flag coimmunoprecipitated with MBP-GEF14 (PD: α-MBP/MBP-GEF14; WB: α-Flag). Middle shows the expression of ROP6-Flag proteins (WB: α-Flag for input control), and Bottom showed the input recombinant proteins (WB: α-MBP, for input control).

**Supplemental Figure 7.**
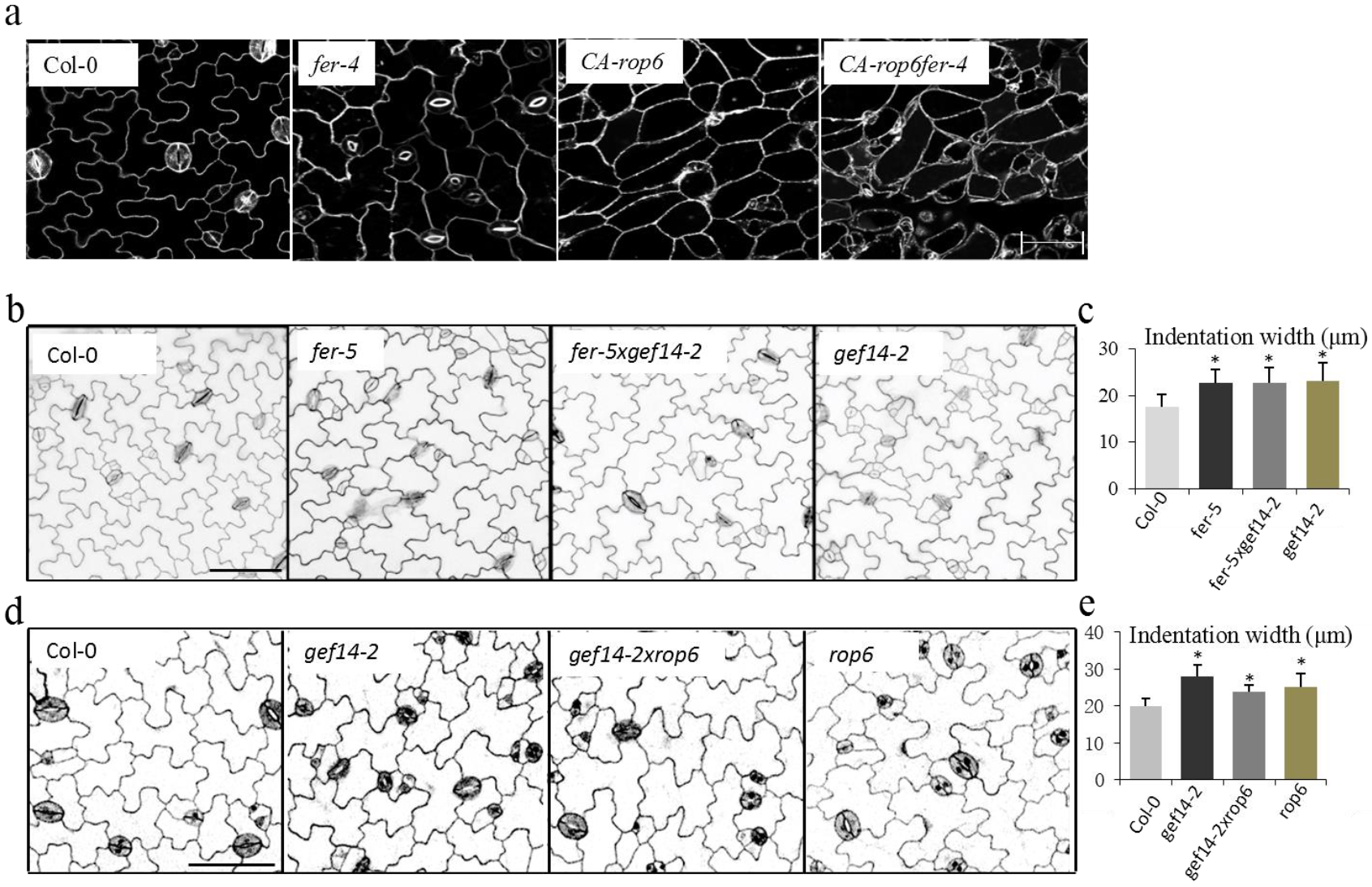
FER, GEF14, and ROP6 genetically act in the same signaling pathway in regulating PC morphogenesis. **a** ROP6 genetically acts in the same pathway as FER in regulating PC morphogenesis. PC morphogenesis was characterized by wild-type, *fer-4, CA-ROP6*, and *CA-ROP6fer-4*. Scale bar, 50 μ m. **b and c**, GEF14 genetically acts in the same signaling pathway as FER in regulating PC morphogenesis. PC phenotypes were characterized by 2 DAG wild-type, *gef14-2, fer-4* and *fer-4gef14-2* mutants (b). The degree of PC interdigitation was determined by the width of indentation (c). **d and e**, ROP6 genetically acts in the same signaling pathway as GFE14 in regulating PC morphogenesis. PC phenotypes were characterized by 2 DAG wild-type, *gef14-2, rop6* and *gef14-2rop6* mutants (d). The degree of 2 DAG PC interdigitation was determined by the width of indentation (e). Asterisks indicate the significant difference (p≤0.05, Student’s *t*-test) between the wild type and the mutants in above assays.

